# Cryo-EM analyses of wild-type and oncogenic KIT mutants reveal structural oncogenic plasticity and a novel “Achilles heel” for therapeutic intervention

**DOI:** 10.1101/2022.09.08.506998

**Authors:** Stefan G. Krimmer, Nicole Bertoletti, Yoshihisa Suzuki, Luka Katic, Jyotidarsini Mohanty, Sheng Shu, Sangwon Lee, Irit Lax, Wei Mi, Joseph Schlessinger

## Abstract

The receptor tyrosine kinase KIT and its ligand SCF are required for the development of hematopoietic stem cells, germ cells, and other cells. A variety of human cancers, such as acute myeloid leukemia and mast cell leukemia, are driven by somatic gain-of-function KIT mutations. Here, we report cryo-EM structural analyses of full-length wild-type and two oncogenic KIT mutants, which show that the symmetric arrangement of ligand-occupied KIT dimers is converted into asymmetric D5 homotypic contacts juxtaposing the plasma membrane. Mutational analysis of KIT reveals in D5 region an “Achilles heel” for therapeutic intervention. A ligand-sensitized oncogenic KIT mutant exhibits a more comprehensive and stable D5 asymmetric conformation. A constitutively active ligand-independent oncogenic KIT mutant adopts a V-shaped conformation solely held by D5-mediated contacts. SCF binding to this mutant fully restores the conformation of wild-type KIT dimers, revealing an unexpected structural plasticity of oncogenic mutants that may offer new therapeutic modality.

## Introduction

Over the last decade, valuable insights have been gained into the mechanism of action of receptor tyrosine kinase (RTK) proteins stimulated by their physiological ligands or by aberrantly activated somatic or germ-line disease-causing mutations (1). New molecular mechanisms were revealed by structural and biochemical analyses combined with experiments in which the activities of mutant RTK proteins expressed in cultured cells were analyzed and compared. Most of the new mechanistic insights were obtained from structural and biochemical analyses of free or ligand-occupied soluble extracellular domains (ECDs) lacking the transmembrane (TM) and cytoplasmic regions of the RTKs. Similarly, soluble fragments of the cytoplasmic regions of RTKs composed of inactive or phosphorylated tyrosine kinase domains mostly devoid of other cytoplasmic regions were also structurally and biochemically explored.

The first images of detergent-solubilized preparations of ligand-occupied full-length RTK dimers were obtained by analyzing negatively-stained EM samples of full-length EGF-induced EGF receptor (EGFR) dimers (2), SCF-induced KIT dimers (3), and PDGF-induced PDGF receptor (PDGFR) dimers (4). These studies revealed low-resolution structures of the ligand-occupied ECDs of these RTKs that are consistent and in accord with the high-resolution structures of corresponding ligand-occupied ECD complexes determined by X-ray crystallography.

Several reports have recently described high-resolution cryo-EM analyses of full-length RTKs solubilized in detergent, amphipol or nanodiscs, including insulin-occupied insulin receptor (5), EGF or TGF-α-occupied EGFR (6), and heterodimeric complexes of full-length ErbB2 with ErbB3 (7). These structural analyses provide valuable insights into the stoichiometry of insulin binding to insulin receptor, reveal a role for the membrane-proximal tips of the ECD in ligand-induced EGFR activation, and provide insight into the mechanism of ErbB2 and ErbB3 heterodimerization within the framework of full-length structures of members of the EGFR family. However, neither the transmembrane domains nor the cytoplasmic regions were resolved in the cryo-EM structures of these full-length receptors, probably due to the dynamic nature and flexibility of the cytoplasmic domains relative to the rest of the RTK molecules.

Genetic studies demonstrated that KIT and its ligand SCF (stem cell factor) are essential for development of hematopoietic stem cells, germ cells, melanocytes, and Cajal cells of the gastrointestinal tract (8, 9). Moreover, gain-of-function KIT mutations were identified in various human cancers and shown to function as critical oncogenic drivers of different cancers including acute myeloid leukemia (AML), mast cell leukemia (MCL), gastrointestinal stromal tumors as well as melanoma (10, 11).

KIT is a member of the type-III subfamily of RTKs, which also includes PDGFR*α*, PDGFR*β*, Csf1R (colony stimulating factor 1 receptor or FMS), and Flt3R (FMS-like tyrosine kinase 3 receptor) (12). Detailed biochemical and structural analyses of KIT have established a common activation mechanism of the type-III RTK subfamily (13–17). Binding of pre-existing SCF dimers induces KIT dimerization, enabling homotypic contacts between two membrane-proximal domains D4 and D5 of KIT. These homotypic contacts enable *trans* autophosphorylation, stimulation of tyrosine kinase activity, and recruitment and activation of multiple intracellular signaling pathways essential for mediating KIT’s pleiotropic response. Similar mechanisms of activation and cell signaling were revealed for other members of the type-III RTK subfamily (18).

In this report, we describe cryo-EM analyses of the structures of full-length ligand-induced KIT dimers. The cryo-EM structures of ligand-induced wild-type and ligand-sensitized oncogenic KIT mutants provide mechanistic insights into a new conserved “Achilles heel” in D5 of KIT, a hotspot for activating somatic mutations (8, 18), which can be therapeutically targeted for inhibition of oncogenically activated KIT and other type-III RTKs. The structural analyses demonstrate that the 2-fold symmetric arrangement of SCF bound to the KIT ligand-binding region (D1–D3) and of the salt bridges mediating homotypic D4:D4’ contacts is converted into an asymmetric arrangement of D5:D5’ contacts. The asymmetric arrangement is caused by a shift of opposing interfaces by one residue relative to each other, resulting in a different tilt of the two interacting D5 regions that juxtapose the cell membrane. Comparison of the cryo-EM structures of ligand-induced full-length KIT dimers to those of an oncogenic ligand-sensitized KIT mutant and of a constitutively active (ligand-independent) oncogenic KIT mutant provides important information about a molecular mechanism that governs KIT-driven oncogenic transformation. These structural analyses also reveal an unexpected plasticity in the structure and activity of the constitutively activated oncogenic KIT mutant that primarily acts intracellularly.

## Results

To gain further insights into the physiological and oncogenic activation mechanism of KIT, we pursued structural investigations of SCF-induced full-length KIT dimers and of oncogenically activated KIT mutants using cryo-EM. We expressed wild-type and oncogenic mutants of full-length KIT in ExpiSF9 insect cells. To improve protein yield, a K623A mutation, inactivating the tyrosine kinase activity of KIT, was introduced into all constructs used for structural studies. SCF-bound or free KIT preparations were solubilized in a detergent, followed by reconstitution in an amphipol. Reconstitution of the complexes in an amphipol was necessary to achieve a uniform particle distribution in the vitreous ice of the holey cryo-EM grids. After reconstitution, the complexes were covalently crosslinked with glutaraldehyde during purification by density gradient ultracentrifugation (19). This procedure was necessary to prevent dissociation of the labile complexes.

We collected four cryo-EM datasets of wild-type and mutants of full-length KIT (Table S1): SCF-occupied wild-type KIT, SCF-occupied KIT(DupA502,Y503) designated ligand-sensitized oncogenic KIT mutant, free KIT(T417I,Δ418-419) designated constitutively active oncogenic KIT mutant, and SCF-occupied KIT(T417I,Δ418-419). In all four cryo-EM maps, only the extracellular regions of the complexes were resolved. Despite extensive attempts, we were not able to focus particle alignment on the dimeric cytoplasmic region (Fig. 1). We surmise that the inability to obtain a cryo-EM map of the dimeric cytoplasmic domains of wild-type KIT and its oncogenic mutants may arise from the small size of the cytoplasmic domain (95 kDa), and primarily from the dynamic nature of the cytoplasmic region that enables the tyrosine kinase domain of KIT to mediate efficient *trans* autophosphorylation within the context of a dimeric receptor complex. All recently published cryo-EM analyses of full-length RTKs similarly reported an inability to resolve structures of the cytoplasmic domain of ligand-activated full-length RTKs (5–7, 20). This suggests that structural flexibility of the linker connecting the ECD with the transmembrane and cytoplasmic domains is a common feature of RTKs.

**Fig. 1.**
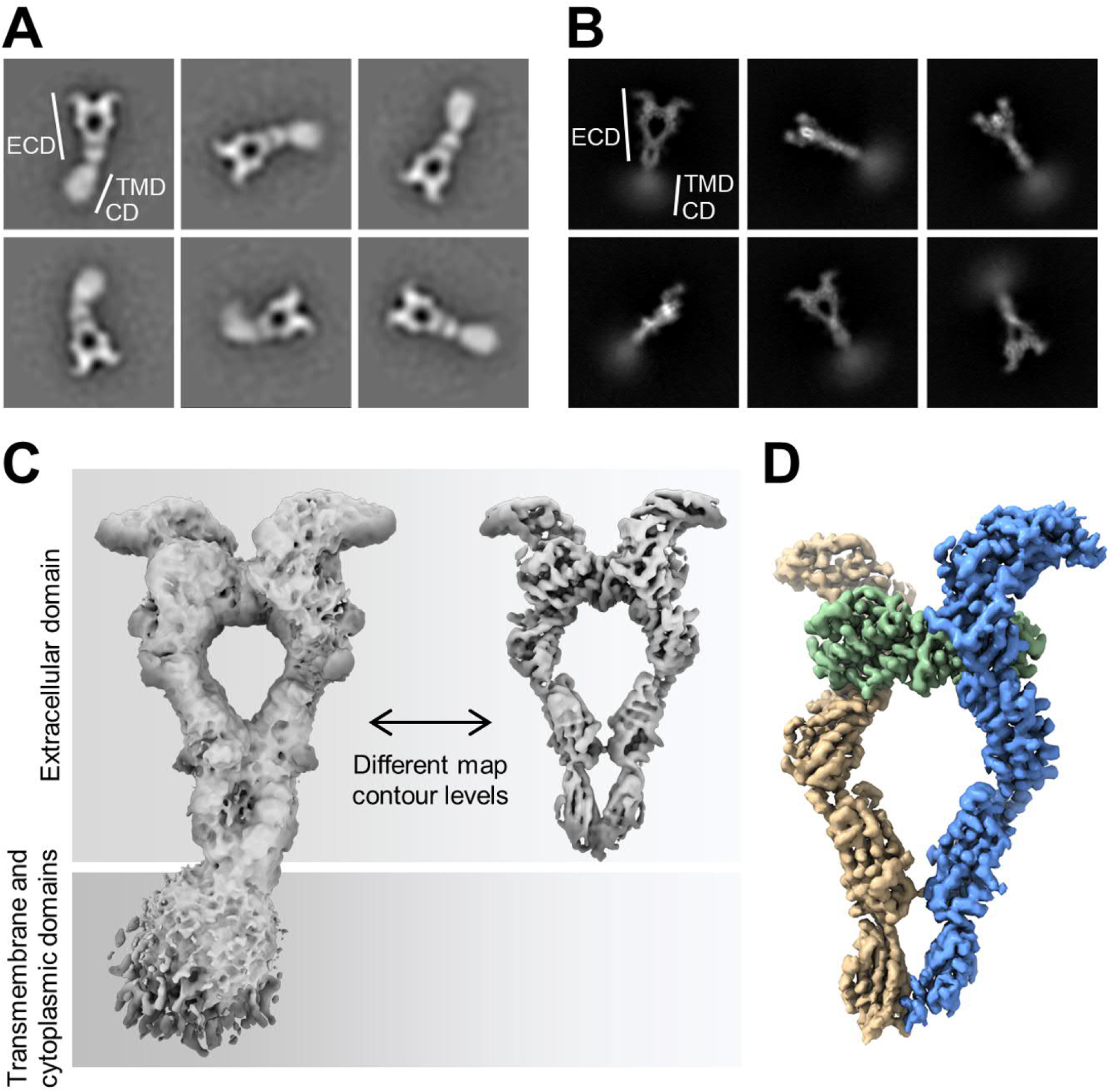
Cryo-EM analysis of full-length wild-type KIT:SCF dimers reconstituted in an amphipol environment resolves the extracellular domain to high resolution. (*A*) Representative negative staining EM 2D class averages. Regions corresponding to the extracellular domain (ECD), the transmembrane domain (TMD), and cytoplasmic domain (CD) are indicated. (*B*) Representative cryo-EM 2D class averages (front and side views only). The extracellular domain is sharply defined, whereas the cytoplasmic domain is blurred out. (*C*) Cryo-EM 3D reconstruction after global refinement of the entire KIT:SCF complex. The cryo-EM map is displayed at low (left) and high (right) map contour levels. (*D*) Cryo-EM 3D reconstruction of the ECD of KIT:SCF dimers after local ECD refinement and post-processing (overall resolution: 3.45 Å; KIT protomer A in beige, KIT protomer B in blue, SCF homodimer in green).

### The symmetric arrangement of ligand-occupied KIT dimers is converted into asymmetric D5:D5’ contacts juxtaposing the plasma membrane

We determined the cryo-EM map of the ECD of full-length wild-type KIT:SCF at a global resolution of 3.45 Å (Fig. 2 and S1; local resolution map in Fig. S1B). 3D variability analysis (21) of the cryo-EM map (Fig. S1G) reveals that the complex between neighboring domains D5 and D5’ (D5:D5’) is oscillating perpendicularly relative to the view of Fig. 2A, explaining the lower resolution of D5:D5’ relative to rest of the ECD (Fig. S1B). The conformation of SCF bound to the ligand-binding region of KIT (D1–D3) is virtually identical to the conformation in the crystal structure of SCF in complex with the soluble ECD of KIT (17). Furthermore, the conformations of domains D1–D4 and D1’–D4’ in the cryo-EM structure are very similar to their conformations in the crystal structure (Fig. S2). The cryo-EM map also clearly reveals the salt bridge between E386 and R381 responsible for mediating the highly conserved D4:D4’ interface (Fig. S1E; (17)). By contrast, the cryo-EM structure reveals a novel asymmetric interface formed between two neighboring domains D5 and D5’ with a buried surface area of 292 Å^2^ (Fig. 2B). This D5:D5’ interface is not present in the crystal structure of the soluble ECD (Fig. S2B). It is clear that direct D5:D5’ contact formation depends upon the integrity of the KIT receptor, requiring the presence of transmembrane and cytoplasmic domains in full-length KIT. D5 of KIT exhibits an Ig-like β-sandwich fold composed of anti-parallel β-sheets βA-βB-βE-βD and βC-βF-βG. We refrained from including β-strands βD and βD’, and the loops proximal to the membrane into the model of D5:D5’ (Fig. S2B) due to limited resolution of these regions in the cryo-EM map. In comparison to the uncomplexed conformation in the crystal structure, formation of the D5:D5’ interface requires Y418 and Y418’ to undergo a conformational change from a solvent-exposed to a core-buried conformation (Fig. S3). Superposition of the two KIT protomers reveals that domain D5’ is strongly tilted toward domain D5, resulting in an asymmetric conformation of D5:D5’ (Fig. S2A–C). The asymmetric conformation of D5:D5’ is enabled by the flexibility of the linkers connecting D4 to D5 and D4’ to D5’, whereas the overall tertiary conformation of the two domains of D5:D5’ remains largely unaffected (Fig. S2D and S2E). In contrast to D5:D5’, the remaining part of the ECD of the KIT:SCF cryo-EM structure exhibits a 2-fold symmetric conformation (Fig. S2A and S2C).

**Fig. 2.**
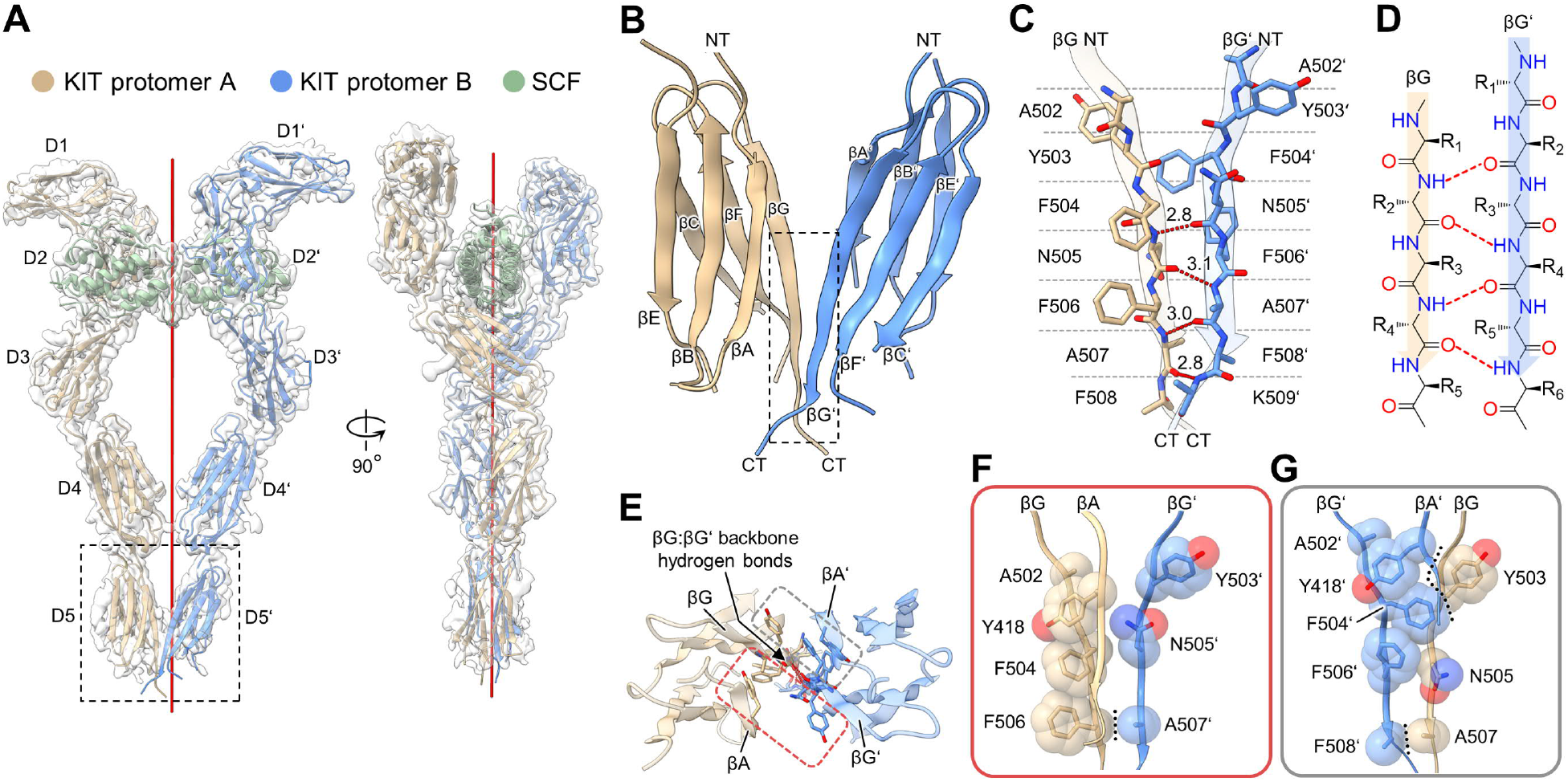
Cryo-EM structure of full-length wild-type KIT:SCF dimers reveals an asymmetric conformation of D5:D5’ contacts. (*A*) Cryo-EM map (gray volume) of the ECD of KIT:SCF dimers. The structural model fitted into the cryo-EM map shows KIT protomer A in beige, KIT protomer B in blue, and SCF homodimer in green. The red line represents the 2-fold rotation symmetry axis. The same color-code is used in *A*–*G*. (*B*) Close-up view of the boxed asymmetric D5:D5’ complex from *A*. NT, N-terminus; CT, C-terminus. (*C*) Close-up view of the boxed asymmetric βG:βG’ interface from *B*. Backbone hydrogen bonds are shown as red dotted lines (distances labeled in Å). (*D*) Schematic representation of the asymmetric βG:βG’ backbone interactions. Relative to β-strand βG, β-strand βG’ is shifted by one residue toward the N-terminus and rotated by 180°. Backbone hydrogen bonds are shown as red dotted lines. (*E*–*G*) Side chain interactions at the D5:D5’ interface. (*E*) Top view of D5:D5’ indicating side chain interactions at site-I (red box) and site-II (gray box). Residues with side chains participating in interface interactions are shown as sticks. (*F*–*G*) Side chain interactions at site-I (*F*) and site-II (*G*) of the D5:D5’ interface. Hydrophobic packing interactions at the interface are delineated by black dotted lines. Side chains are shown as sticks, and their van der Waals radii are shown as semitransparent spheres to highlight shape complementarity. Backbone atoms are omitted for clarity. The side chain of F508’ was omitted from the model due to poorly defined cryo-EM density.

Residues that take part in formation of the D5:D5’ interface are well defined in the cryo-EM map (Fig. S1F). The D5:D5’ interface is mainly formed between two neighboring β-strands βG and βG’, which are running in the same direction, via asymmetric interactions (Fig. 2C and 2D). Relative to βG, the residues of βG’ are shifted by a single amino acid residue toward the N-terminus. The resulting arrangement enables the formation of four backbone hydrogen bonds between βG:βG’. The backbone interactions of βG:βG’ divide the side chain interactions at the interface into two separated sites, which we designate as site-I and site-II (Fig. 2E–G). All side chains at the interface are hydrophobic, except for the hydrophilic side chains of N505 and N505’, which are solvent exposed. Because N505’ is located at the center of site-I, the hydrophilic side chain of N505’ largely prevents hydrophobic interactions at the interface, allowing only hydrophobic interactions between F506 and A507’ (Fig. 2F, dotted line). In contrast to its location in site-I, N505 in site-II is shifted by one residue toward the C-terminus due to the asymmetric interface, enabling hydrophobic interactions between Y503 and F504’ (Fig. 2G). Furthermore, the shift of N505 enables Y418’ in βA’ to move closer toward the interface, enabling hydrophobic interactions between Y418’ and Y503.

### Integrity of the D5:D5’ interface is essential for ligand-induced KIT activation

To explore the role of the D5:D5’ interface in SCF-induced KIT activation, we mutated residues involved in mediating hydrophobic side chain interactions at the asymmetric interface and evaluated their impact on tyrosine kinase activation of KIT upon stimulation with SCF in cells expressing wild-type or KIT mutants. We selected F504 and F506 for mutational studies because these hydrophobic residues are conserved in all type-III RTK family members (Fig. 3A). We further tested whether simultaneous disruption of D4:D4’ and D5:D5’ interfaces may have a synergistic effect on impairing KIT activation. Accordingly, we explored the impact of mutations in the D5:D5’ interface alone or in combination with R381A, a mutation disrupting the salt bridge between R381 and E386 at the D4:D4’ interface (Fig. S1E). In total, the properties of seven KIT mutants—KIT(R381A), KIT(F504A), KIT(F506A), KIT(F504A,F506A), KIT(R381A,F504A), KIT(R381A,F506A), and KIT(R381A,F504A,F506A)—were analyzed and compared. Initial expression experiments showed that KIT(F504A,F506A) and triple mutant KIT(R381A,F504A,F506A) are not expressed on the cell surface (Fig. S4). Therefore, these two mutants were excluded from further analysis.

**Fig. 3.**
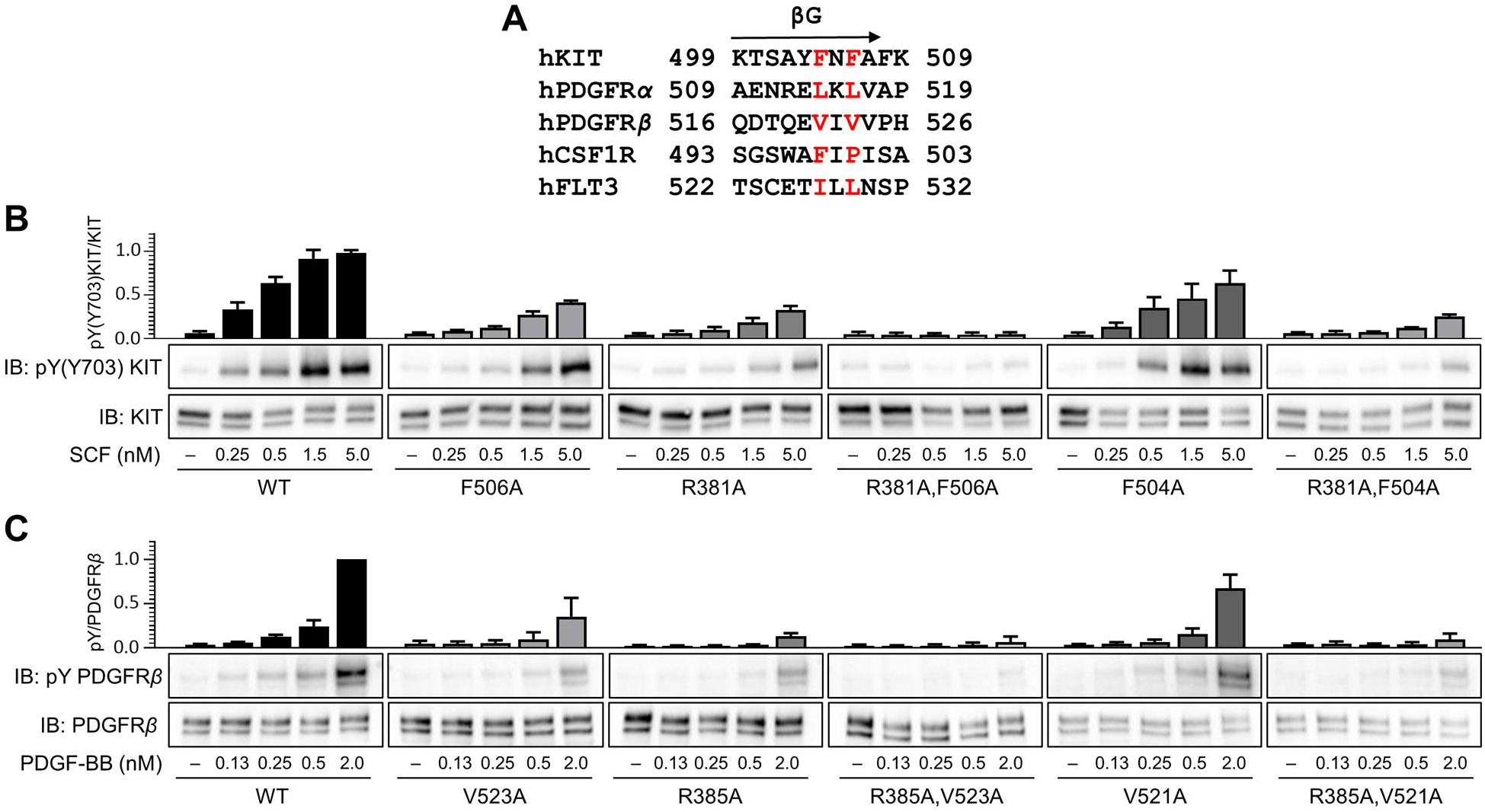
Tyrosine autophosphorylation of wild-type and mutants of KIT and PDGFR*β* in response to ligand stimulation. (*A*) Structure-based sequence alignments of β-strands βG of human type-III RTK family members. Functionally conserved residues are highlighted in red. (*B*) Lysates from SCF stimulated or unstimulated NIH 3T3 cells stably expressing wild-type KIT or KIT mutants were subjected to immunoprecipitation with anti-KIT antibody followed by SDS-PAGE and immunoblotting with either anti-phospho-KIT(Y703) or anti-KIT antibodies. Tyrosine autophosphorylation of wild-type or KIT mutants in response to different concentrations of SCF was monitored. Representative blots are shown from experiments performed in triplicate. Quantification of KIT(Y703) phosphorylation is normalized relative to KIT expression levels. Quantification results show mean ±s.d. of three independent experiments. (*C*) PDGF-BB stimulated or unstimulated MEF cells stably expressing wild-type HA tagged PDGFR*β* or PDGFR*β* mutants were lysed and subjected to immunoprecipitation with anti-HA antibody followed by SDS-PAGE and immunoblotting with anti-PDGFR*β* antibody or anti-phosphotyrosine antibody. Representative blots are shown from experiments performed in triplicate. Quantification of PDGFR*β* phosphorylation is normalized relative to PDGFR*β* expression levels. Quantification results show mean ±s.d. of three independent experiments.

We next compared the ability of SCF to stimulate cultured NIH 3T3 cells stably expressing wild-type or KIT mutants, matched for expression level, to induce tyrosine autophosphorylation of KIT. The experiment presented in Fig. 3B shows that the mutant KIT molecules exhibit either a partial or full loss of SCF-induced tyrosine autophosphorylation compared to SCF-induced tyrosine autophosphorylation of wild-type KIT expressed in these cells. The single mutations F504A and F506A showed reduced ligand-induced KIT activation with the F506A mutant of KIT exhibiting approximately 50% reduced tyrosine autophosphorylation when the cells were stimulated with 1.5 nM of SCF, a saturating ligand concentration for wild-type KIT expressed in these cells. Notably, double mutation R381A,F506A exhibited a strong synergistic inhibition manifested by complete loss of SCF-induced tyrosine autophosphorylation of KIT. Double mutation R381A,F504A, on the other hand, showed less synergistic inhibition on SCF-induced tyrosine autophosphorylation of KIT. Taken together, these experiments reveal a second “Achilles heel” for ligand activation in D5 of KIT in addition to the “Achilles heel” previously identified in D4 of KIT (17).

We have previously demonstrated that R385A mutation in PDGF receptor *β* (PDGFR*β)* disrupts the formation of salt bridges mediating D4:D4’ homotypic contacts that are essential for PDGF-induced PDGFR*β* activation (16). Structure-guided sequence alignment of β-strand βG of KIT with βG of other type-III RTK family members shows conservation of hydrophobic amino acids corresponding to F504 and F506 in KIT (Fig. 3A). The experiment presented in Fig. 3C shows that cells expressing V521A or V523A mutants (corresponding to KIT mutants F504A and F506A) exhibited reduced PDGF-induced tyrosine autophosphorylation of PDGFR*β* compared to cells matched for expression of wild-type PDGFR*β*. Moreover, PDGF stimulation of PDGFR*β* harboring V521A or V523A mutations in combination with a R385A mutation that disrupts D4:D4’ homotypic contacts resulted in synergistic inhibition of PDGFR*β* activation (Fig. 3C). This experiment suggests that PDGFR*β* and perhaps other type-III RTK members may utilize a similar mechanism of receptor activation through ligand-induced formation of asymmetric D5:D5’ homotypic associations.

### The ligand-sensitized oncogenic KIT mutant exhibits a more comprehensive and stable D5:D5’ interface

To explore the structural mechanism of the ligand-sensitizing effect of mutation DupA502,Y503 of KIT, we determined the cryo-EM map of the extracellular region of full-length KIT(DupA502,Y503):SCF dimers at an overall resolution of 3.17 Å (Fig. S5; local resolution map in Fig. S5B). The conformations of domains D1–D4, D1’–D4’, and bound SCF are virtually identical to their conformations in the cryo-EM structure of wild-type KIT:SCF dimers. However, a more comprehensive asymmetric D5:D5’ interface than in wild-type KIT was resolved in the structure of KIT DupA502,Y503 mutant (Fig. 4), revealing a 64% larger buried surface area (479 Å^2^). Additional D5:D5’ contacts are caused by introduction of residues DupA502, DupA502’, DupY503, and DupY503’ at the βG:βG’ interface. Furthermore, introduction of these residues also shifts N505’ by two residue positions toward the C-terminus of the molecule (Fig. 4B). Occupying the former position of N505’, DupY503’ now forms tight hydrophobic interactions with Y418 in βA of β-sheet βA-βB-βE, contributing to the compact conformation of the D5:D5’ interface. By contrast, in wild-type KIT:SCF dimers the polar side chain of N505’ repulses Y418 in βA, resulting in less extensive D5:D5’ contacts (Fig. 4B). In contrast to the conformational change of β-sheet βA-βB-βE, there is no significant conformational change of β-sheet βA’-βB’-βE’ compared to wild-type KIT:SCF (Fig. 4C). In wild-type KIT:SCF, N505 is distant from and therefore does not make a direct contact with Y418’ in βA’ because of the asymmetric βG:βG’ interface conformation (Fig. 4D). Therefore, the insertion of DupA502 and DupY503 and the resulting shift of N505 by two residue positions to the C-terminus does not affect the conformation of β-sheet βA’-βB’-βE’.

**Fig. 4.**
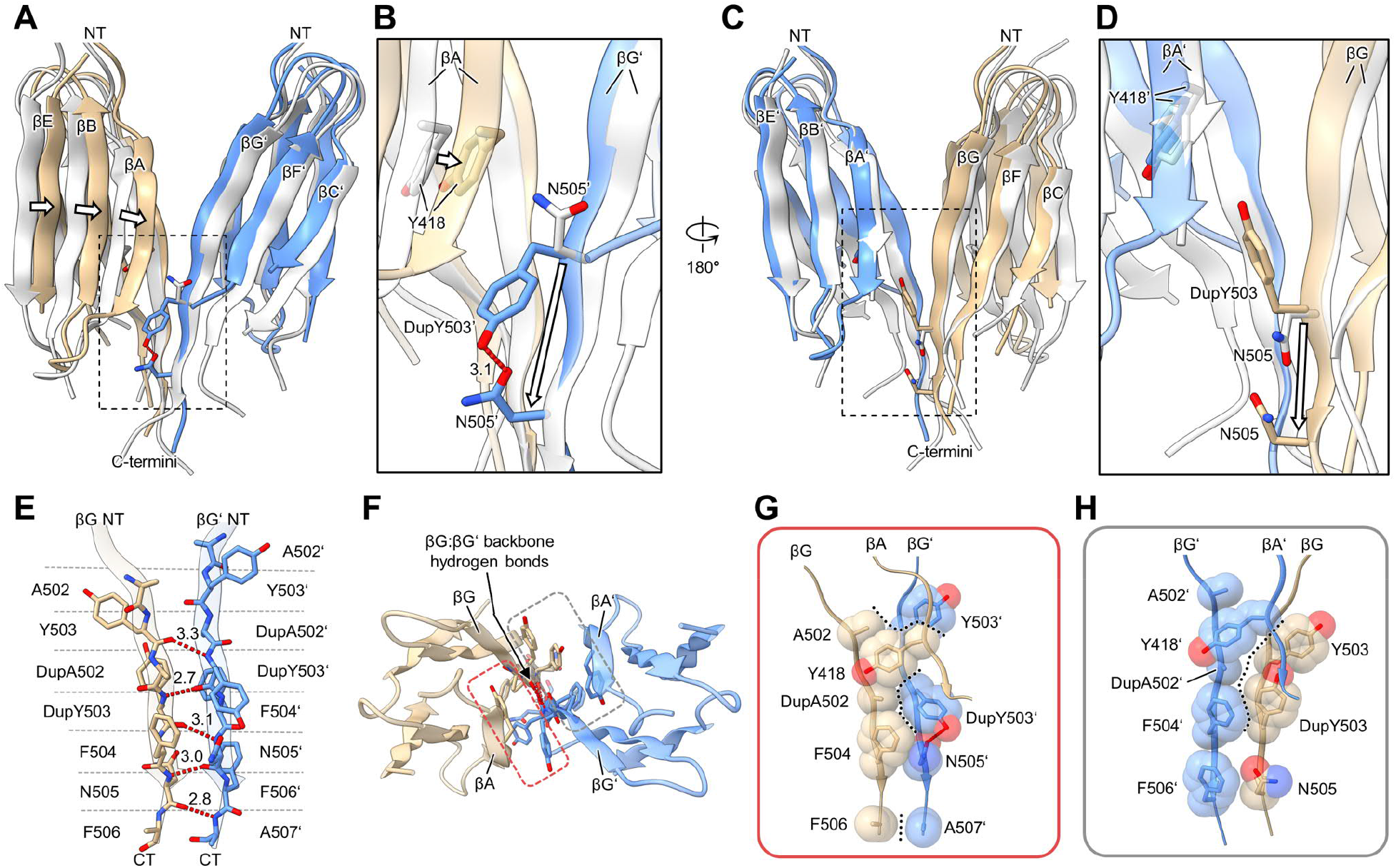
Cryo-EM structure of full-length KIT(DupA502,Y503):SCF dimers reveals the ligand-sensitizing mechanism of the oncogenic KIT mutant. (*A*) Superposition of D5:D5’ of KIT(DupA502,Y503):SCF dimers (D5 in beige, D5’ in blue) and D5:D5’ of wild-type KIT:SCF dimers (D5 and D5’ both in white). The conformational change of β-sheet βA-βB-βE is indicated by black outlined arrows. Structures were aligned on their ECDs excluding D5:D5’ from the calculation of the alignment. The same color-code is used in *A*–*H*. NT, N-terminus. (*B*) Enlarged view of boxed residues Y418, DupY503’, and N505’ from *A*. The shift in the location of N505’ between wild-type and DupA502,Y503 mutant is indicated by the black outlined arrow. (*C*) Same superposition as in *A*, rotated by 180°. (*D*) Enlarged view of boxed residues Y418’, DupY503, and N505 from *C*. The shift in the location of N505 between wild-type and DupA502,Y503 mutant is indicated by the black outlined arrow. (*E*) Asymmetric D5:D5’ interface formed by β-strands βG and βG’. Red dotted lines indicate backbone hydrogen bonds (distances labeled in Å). (*F*–*H*) Side chain interactions at the D5:D5’ interface. (*F*) Top view of D5:D5’ indicating side chain interactions at site-I (red box) and site-II (gray box). Resides with side chains participating in interface interactions are shown as sticks. (*G*–*H*) Side chain interactions at site-I (*G*) and site-II (*H*) of the D5:D5’ interface. Hydrophobic packing interactions at the interface are delineated by black dotted lines. Side chains are shown as sticks, and their van der Waals radii are shown as semi-transparent spheres to highlight shape complementarity. Backbone atoms are omitted for clarity. The side chain of F506 was omitted from the model due to poorly defined cryo-EM density.

Similar to the D5:D5’ interface of wild-type KIT:SCF dimers, the D5:D5’ interface of KIT(DupA502,Y503):SCF dimers has an asymmetric conformation (Fig. 4E). At the interface, residues of β-strand βG’ are shifted by one residue toward the N-terminus relative to residues of β-strand βG. Whereas four hydrogen bonds are formed at the βG:βG’ interface of wild-type KIT:SCF dimers (Fig. 2C), the more extensive contacts in D5:D5’ of KIT(DupA502,Y503):SCF dimers result in formation of five backbone hydrogen bonds at the βG:βG’ interface (Fig. 4E).

Compared to D5:D5’ hydrophobic interface interactions in wild-type KIT:SCF dimers (Fig. 2E–G), the D5:D5’ interface of KIT(DupA502,Y503):SCF dimers experiences a significant increase in hydrophobic interface interactions (Fig. 4F–H). This is due to insertion of DupY503 and DupY503’ at the interface. Whereas the hydroxyl-groups of DupY503 and DupY503’ are solvent-exposed, their aromatic rings are involved in hydrophobic contacts. In addition to the insertion of the hydrophobic residues, N505 and N505’ are C-terminally shifted by two residues, which prevents the disruption of hydrophobic interactions at the interface.

### The extracellular domain of the constitutively active oncogenic KIT mutant adopts a V-shaped conformation

We further used cryo-EM to determine the structure of full-length KIT(T417I,Δ418-419)—a constitutively active KIT mutant—in the absence of SCF. Negative staining EM and cryo-EM 2D class averages reveal the formation of KIT(T417I,Δ418-419) dimers even in the absence of SCF binding (Fig. S6A and S6B). As indicated by the blurred-out regions in the 2D class averages, the dimeric complex of KIT(T417I,Δ418-419) has a high conformational flexibility which is due to lack of stabilization caused by SCF binding to the ligand-binding region. Because of the high conformational flexibility, we were unable to achieve a high-resolution 3D reconstruction of this cryo-EM dataset. Therefore, we refined two different cryo-EM maps at medium to low resolution. The first cryo-EM map is a global refinement of the entire particle without the application of a mask (Fig. S6C). This map is relatively well resolved for D4D5 and D4’D5’, revealing a V-shaped conformation for them (Fig. 5A and 5B). The cryo-EM map features two characteristic bulges at a map region corresponding to the D5:D5’ complex. We docked the crystal structure of the D4D5 fragment dimer of KIT(T417I,Δ418-419) into the map and achieved an overall good fit. The angle between the two protomers in the crystal structure appears slightly larger than the angle in the cryo-EM map, which can be attributed to the high degree of structural flexibility of this complex. The docking pose of the D4D5 fragment dimer explains the two characteristic bulges in the map with the tilted orientation of the two domains of D5:D5’.

**Fig. 5.**
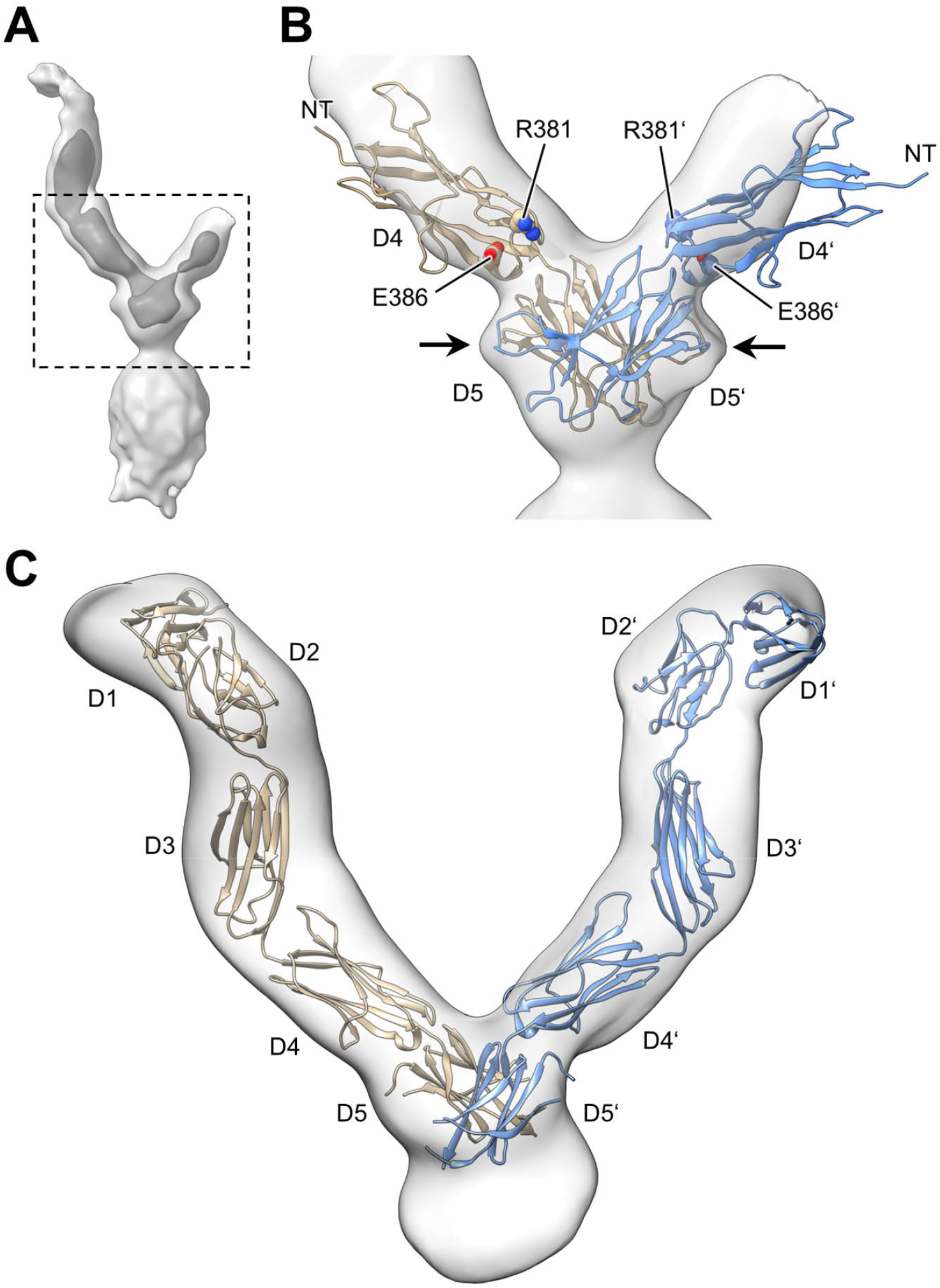
Cryo-EM map of full-length KIT(T417I,Δ418-419) dimers. (*A*) Cryo-EM map of a global refinement of the entire complex. The cryo-EM map is displayed at low (light gray) and high (dark gray) contour levels. (*B*) Close-up view of the boxed region from *A*. The crystal structure of the D4D5 fragment dimer of KIT(T417I,Δ418-419) (PDB ID 4PGZ) was rigid-body docked into the cryo-EM map. Fragment D4D5 of protomer A in beige, fragment D4’D5’ of protomer B in blue. Black arrows indicate the two bulges in the cryo-EM map resulting from the tilted conformation of D5:D5’. NT, N-terminus. (*C*) Cryo-EM map of a masked refinement of the ECD. Two copies of the crystal structure of monomeric KIT (PDB ID 2EC8) were docked into the map.

We refined the second cryo-EM map of the dataset as a masked refinement of the ECD, with the exclusion of the cytoplasmic domain (Fig. S6D). In this low-resolution map, the extracellular domains D1–D5 and D1’–D5’ are fully resolved, exhibiting a characteristic V-shaped conformation (Fig. 5C). The resolution of the map enables visualizing individual domains as separate density blobs at high contour levels. Rigid-body docking of two copies of the previously reported crystal structure of the monomeric ECD of KIT into the map followed by slight manual adjustment of D5 resulted in a good fit. The resulting model shows the ECDs adopting a V-shaped conformation, with a similar angle as in the crystal structure of the D4D5 fragment dimer of KIT(T417I,Δ418-419) (Fig. S6E). The only interactions between the ECDs are mediated via their D5:D5’ contacts (the ‘bottom’ of the V), whereas D1–D4 and D1’–D4’ are distant from each other (the ‘arms’ of the V).

Consequently, the two cryo-EM maps of full-length KIT(T417I,Δ418-419) dimers confirm the V-shaped conformation previously determined for the crystal structure of D4D5 fragment dimers of KIT(T417I,Δ418-419) (15). Furthermore, the cryo-EM results are fully consistent with and support the molecular interactions at the D5:D5’ interface as seen in the crystal structure of D4D5 fragment dimers of KIT(T417I,Δ418-419).

Compared to wild-type KIT:SCF, mutation T417I,Δ418-419 induces a major conformational change at the D5:D5’ interface (Fig. S7). Whereas the D5:D5’ interface of wild-type KIT:SCF in the cryo-EM structure is asymmetric and mainly formed between βG:βG’, the D5:D5’ interface of KIT(T417I,Δ418-419) dimers in the crystal structure is symmetric and formed between βA:βG’ and βA’:βG. This conformational change is associated with a large increase in D5:D5’ interdomain interactions resulting in a substantial increase of buried surface area (1001 Å^2^) compared to wild-type KIT:SCF (292 Å^2^). The βG:βG’ interface of wild-type KIT:SCF dimers enables parallel orientation of the two domains of D5:D5’, whereas the βA:βG’, βA’:βG interface of KIT(T417I,Δ418-419) dimers induces a strong tilt between the two domains of D5:D5’ (Fig. S7C and S7F). This tilt results in more than doubling of the distance between the SCF binding regions D1–D3 and D1’– D3’ in comparison to wild-type KIT:SCF (Fig. S7A and S7D). Furthermore, whereas in wild-type KIT:SCF dimers the C-terminal ends of domains D5 and D5’ are located next to each other (distance: 4.5 Å, measured between Cα atoms of A507 and F508’, Fig. S7C), in KIT(T417I,Δ418-419) dimers the distance between the C-terminal ends of domains D5 and D5’ are far apart from each other (distance: 15.0 Å, measured between Cα atoms of N505 and N505’, Fig. S7F).

### The wild-type conformation of KIT is restored by ligand binding to the constitutively active oncogenic KIT mutant

Since the V-shaped conformation of the ECD of KIT(T417I,Δ418-419) dimers is vastly different from the conformation of the ECD of wild-type KIT:SCF, we wondered whether SCF is still capable of binding to full-length KIT(T417I,Δ418-419) dimers. Surprisingly, negative staining EM data of SCF-bound KIT(T417IΔ418-419) show the characteristic shape of ligand-bound KIT particles (Fig. S8A), revealing that the SCF homodimer is capable of binding and bringing together the two distant ECDs of the KIT(T417I,Δ418-419) dimer. To gain detailed insight into the mechanism of SCF binding to the oncogenic mutant, we next determined a cryo-EM map of full-length SCF-bound KIT(T417I,Δ418-419) at a global resolution of 3.96 Å (Fig. 6A and S8, local resolution map in Fig. S8C). Overall, the conformations of domains D1–D4, D1’– D4’, and SCF are identical to their conformations in wild-type KIT:SCF complex. The salt bridge between D4:D4’ is clearly resolved in the cryo-EM map (Fig. 6B). Even though the resolution is not sufficient to resolve the interface interactions at a molecular level, the quality of the cryo-EM map of D5:D5’ is sufficient to unambiguously identify an asymmetric D5:D5’ conformation similar to the conformation in wild-type KIT:SCF and interface formation via βG:βG’ (Fig. 6C).

**Fig. 6.**
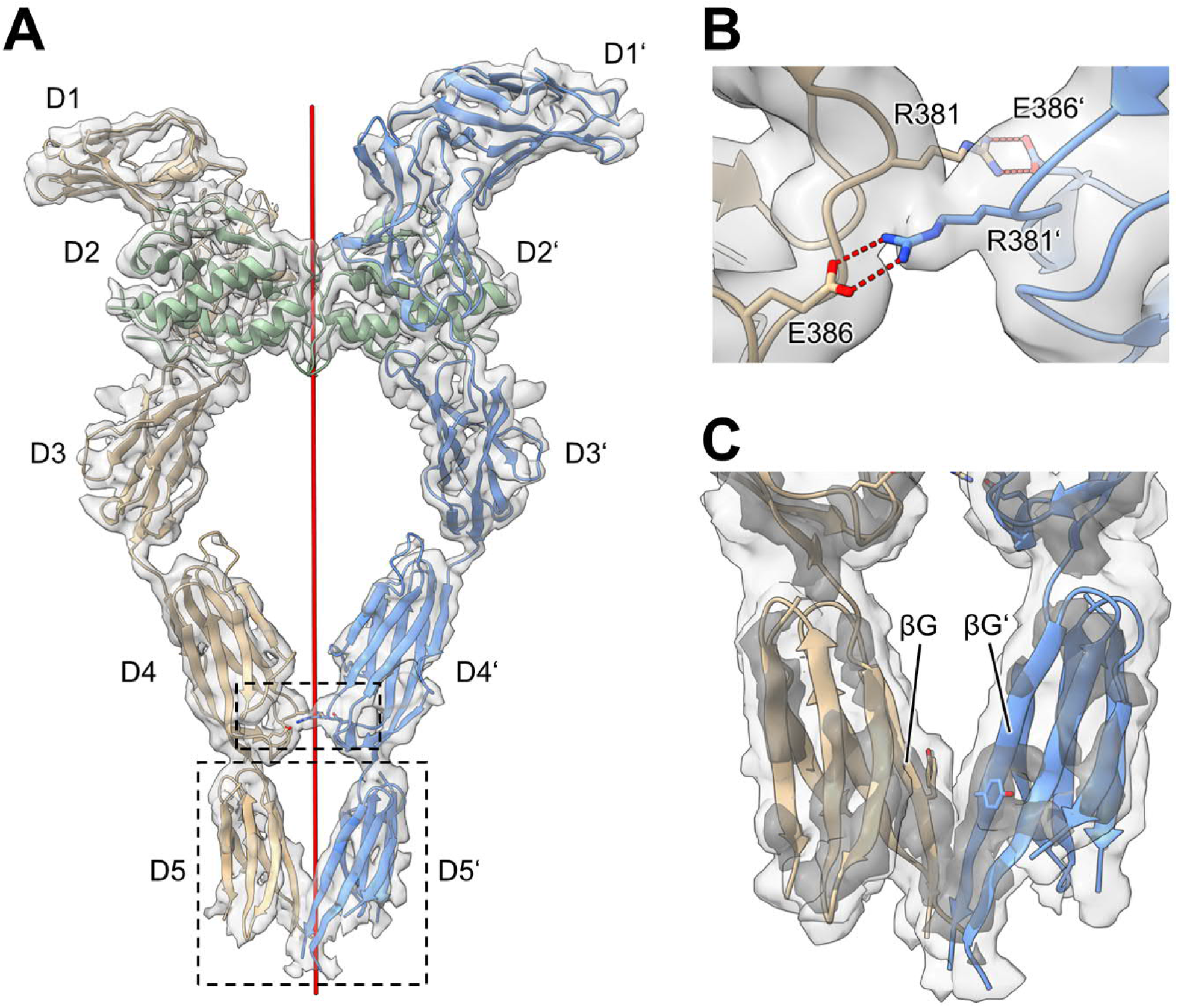
Cryo-EM structure of full-length KIT(T417I,Δ418-419):SCF dimers. (*A*) Cryo-EM map of the ECD of KIT(T417I,Δ418-419):SCF dimers. KIT protomer A in beige, KIT protomer B in blue, SCF homodimer in green. The red line represents the 2-fold rotation symmetry axis. (*B*) Close-up view of the boxed D4:D4’ interface from *A*, detailing the intact homotypic salt bridge interactions between R381 and E386. (*C*) Close-up view of the boxed D5:D5’ complex from *A*. The cryo-EM map is displayed at low (light gray) and high (dark gray) contour levels.

## Discussion

KIT molecules are expressed at the cell membrane as freely diffusing KIT monomers. Binding of SCF dimers to the ligand-binding region (D1–D3) of the extracellular domain of KIT brings two KIT receptors together in the cell membrane. The dramatic increase in local concentration caused by reduced dimensionality of two KIT molecules held together by SCF at the cell membrane combined with the flexibility of inter-domain linkers play an essential role in KIT activation (15). The flexible hinge regions connecting D3 to D4, D4 to D5, and the linker connecting D5 to the transmembrane region enable efficient formation of homotypic D4:D4’ and D5:D5’ contacts as well as additional interactions that may take place between the transmembrane and the cytoplasmic regions with corresponding regions of neighboring KIT molecules. This “zipper-like” mechanism enables formations of D4 and D5 homotypic contacts by weak binding affinities which are not strong enough to mediate dimerization and activation of unoccupied KIT molecules. However, these weak affinities are sufficiently effective due to the high local concentration of KIT at the cell membrane and because of the cooperative action of multiple homotypic contacts in each KIT molecule. The homotypic interactions mediated by D4 and D5 at a region juxtaposing the transmembrane domain set the stage for efficient *trans* autophosphorylation of the tyrosine kinase domain, resulting in stimulation of tyrosine kinase activity followed by recruitment and activation of multiple cellular signaling pathways. A similar “zipper-like” mechanism can be ascribed for the mechanism of activation of other type-III RTK family members stimulated by their specific ligands, and for other RTKs (15).

Our goal in the present study was to use cryo-EM to determine the structures of full-length wild-type KIT and two full-length oncogenic KIT mutants. However, like other recent cryo-EM structural analyses of full-length RTKs, we found out that the dynamic nature of the cytoplasmic region of KIT precludes resolving the structure of the cytoplasmic domain (Fig. 1). Nevertheless, interesting and important new insights were revealed about the structure of the extracellular region in their native full-length state of SCF-occupied KIT dimers, and about the molecular mechanism underlying the action of two different oncogenic KIT mutants.

### Why does the D5:D5’ interface adopt a distinctly asymmetric conformation?

Surprisingly, the cryo-EM structure of ligand-induced full-length wild-type KIT dimers clearly shows that D5:D5’ homotypic contacts adopt an asymmetric conformation, which stands in stark contrast to the symmetric conformation of the remaining part of the ligand-occupied extracellular domain (domains D1–D4, D1’–D4’, and SCF; Fig. S2A and S2C). The asymmetric conformation of the D5:D5’ complex is the result of asymmetric interface interactions, which are mainly formed between the two neighboring β-strands βG and βG’ (Fig. 2B and 2C). The two β-strands βG and βG’ run in the same direction but are rotated by about 180° along the long axis relative to each other. Therefore, to enable the formation of backbone hydrogen bonds, the two β-strands βG and βG’ must engage in asymmetric interactions shifted by one residue (Fig. 2C and 2D).

Two explanations can be proposed for the conversion of a symmetric to an asymmetric conformation from the ligand-binding region to the D5:D5 contacts juxtaposing the TM domain of KIT. One explanation is that the asymmetric conformation of the D5:D5’ contacts juxtaposing the cell membrane may poise the transmembrane and tyrosine kinase domains toward the autophosphorylation reaction that proceeds via an *intermolecular* mechanism. In other words, asymmetric contacts at the extracellular domain close to the cell membrane may set the stage for interactions favoring *trans* autophosphorylation of the cytoplasmic region. An alternative, not mutually exclusive, interpretation is that the asymmetric conformation of D5:D5’ reflects the most energetically stable interface. This interpretation is consistent with the cryo-EM structure of the oncogenic ligand-sensitized KIT(DupA502,Y503) mutant in complex with SCF. This oncogenic mutant exhibits a more comprehensive asymmetric D5:D5’ interface, which significantly increases the buried surface areas to 479 Å^2^ compared to 292 Å^2^ of wild-type KIT. Moreover, the binding affinity of the dimerization reaction of isolated D4D5 fragments of the oncogenic mutant is increased by 10 to 20-fold compared to the corresponding region of wild-type KIT. We favor the interpretation that the asymmetric conformation of D5:D5’ reflects the most energetically stable interface.

### A novel target for therapeutic intervention

To determine the role of D5:D5’ dimerization in the activation process of KIT, we next mutated residues with side chains involved in the formation of the D5:D5’ interface (Fig. 2E–G) and analyzed the tyrosine autophosphorylation activity of KIT upon stimulation with SCF (Fig. 3B). Reducing the hydrophobicity of residues involved in hydrophobic interactions at the D5:D5’ interface compromises the tyrosine autophosphorylation activity of KIT, indicating destabilization of the D5:D5’ interface. Each F506A or F504A mutation reduces the tyrosine autophosphorylation activity of KIT with the F506A mutation showing a stronger inhibition. Double mutant R381A,F506A, which simultaneously disrupts both D4:D4’ and D5:D5’ interfaces, causes a complete loss of tyrosine autophosphorylation activity at any of the tested SCF concentrations (Fig. 3B). Analysis of activities of similar mutations in D5 of PDGFR*β* (Fig. 3C) suggests that formation of asymmetric D5:D5’ contacts may also take place during ligand-induced activation of PDGFR*β* and other members of type-III RTKs. These experiments suggest that pharmacological targeting of D5:D5’ by monoclonal antibodies or other specific binders (protein or non-protein) that occupy D5 and occlude D5:D5’ contact formation may antagonize KIT activation, and thus function as therapeutics for treating cancers or other diseases driven by aberrantly activated or overexpressed KIT proteins. Bi-specific monoclonal antibodies that simultaneously bind to the D4 salt bridge region and to D5 hydrophobic contacts may function as very potent KIT antagonist with broad utility for KIT-driven cancer and other therapeutics. As this mechanism may also play a role in ligand-induced activation of PDGFR*β* and other type-III RTKs, similar pharmacological strategies can be considered for treatment of diseases driven by other activated or overexpressed type-III RTK members.

### Mechanism of activation of the oncogenic ligand-sensitized KIT mutant

The oncogenic KIT(DupA502,Y503) mutant exhibits elevated basal tyrosine kinase activity and can be further activated by SCF binding (15). The tyrosine kinase activity of this mutant relies entirely on the integrity of the salt bridge maintaining the D4:D4’ contacts. The increased dimerization affinity of this mutant shifts the monomer–dimer equilibrium toward dimer formation, thereby disturbing the delicately balanced ligand-mediated activation of wild-type KIT. The elevated basal activity of KIT(DupA502,Y503) is caused by an increased population of active KIT mutant dimers even in the absence of SCF stimulation. Likewise, overactivation upon stimulation with SCF is the result of an increase in the concentration of active KIT(DupA502,Y503):SCF dimers. The cryo-EM structure of KIT(DupA502,Y503):SCF provides a satisfactory explanation for the kinetic properties of this oncogenic mutant. Mutation DupA502,Y503 improves the shape complementarity at the D5:D5’ interface, resulting in additional hydrogen bonds and hydrophobic interactions at the interface (Fig. 4E–H). Increased D5:D5’ interface interactions are responsible for increased affinity and stability of KIT(DupA502,Y503):SCF dimers. Importantly, mutation DupA502,Y503 stabilizes a wild-type-like conformation of D5:D5’, which is compatible with D4:D4’ contact formation and SCF binding. Indeed, introduction of mutation R381A into a background of KIT(DupA502,Y503) abolishes both the elevated basal activity and ligand-stimulation of this oncogenic mutant (15).

### Mechanism of activation of the constitutively active ligand-independent KIT mutant

The oncogenic mutation T417I,Δ418-419 located in D5 of KIT exhibits a constitutively active, ligand-independent tyrosine kinase activity. The tyrosine kinase activity of this mutant is independent of the integrity of the salt bridge mediating D4:D4’ contacts. The binding affinity of isolated D4D5 fragments of KIT(T417I,Δ418-419) dimerization is 200–500 fold higher than the binding affinity of D4D5 fragments of wild-type KIT dimerization (15). Importantly, the majority of KIT(T417I,Δ418-419) molecules are localized intracellularly, and only a small population of this oncogenic mutant is located in the cell membrane (22). The cryo-EM dataset of full-length KIT(T417I,Δ418-419) dimers reveals a V-shaped conformation of the extracellular region. Domains D1–D4 and D1’–D4’ are far apart from each other, whereas D5:D5’ are tightly complexed (Fig. 5). Ligand-independent dimerization of the KIT(T417I,Δ418-419) mutant is entirely mediated by D5:D5’ contacts. The T417I,Δ418-419 mutation of KIT triggers formation of a compact, strongly tilted interface of D5:D5’ causing the V-shaped conformation of KIT(T417I,Δ418-419) dimers (Fig. S7D–F). The tilted D5:D5’ interface is formed by βG:βA’ and βG’:βA, whereas the D5:D5’ interface of wild-type KIT:SCF is only formed by βG:βG’ (Fig. S7C and S7F). As a result, the buried surface area of the D5:D5’ interface increases from 292 Å^2^ to 1001 Å^2^, and thus strongly increases the affinity of homotypic dimerization of KIT(T417I,Δ418-419) compared to wild-type KIT.

### Structural plasticity of an oncogenic KIT mutant

Cryo-EM analysis of the structure of SCF-occupied full-length KIT(T417I,Δ418-419) dimers demonstrates that upon binding of SCF the unique V-shaped conformation of this constitutively activate oncogenic KIT mutant, that is held together solely via stable D5:D5’ contacts, can be entirely converted into a structure nearly identical to the structure of SCF-bound wild-type-like KIT (Fig. 6). Notably, SCF binding fully restores the hallmarks of the conformation of SCF-bound wild-type KIT including restoration of SCF interactions with the D1–D3 regions, restoration of the salt bridges responsible for mediating D4:D4’ contacts and, importantly, restoration of the wild-type conformation of the D5:D5’ interface (Fig. 7). This is quite remarkable because of the large conformational change associated with a major tilt in D5:D5’ contacts that increases the distance between C-terminal amino acids of the two D5 protomers of the oncogenic KIT mutant from 4.5 Å to 15.0 Å. By contrast, the D5:D5’ contacts in SCF-bound KIT(T417I,Δ418-419) dimers are very similar, and likewise asymmetric as those seen in SCF-occupied wild-type KIT dimers.

**Fig. 7.**
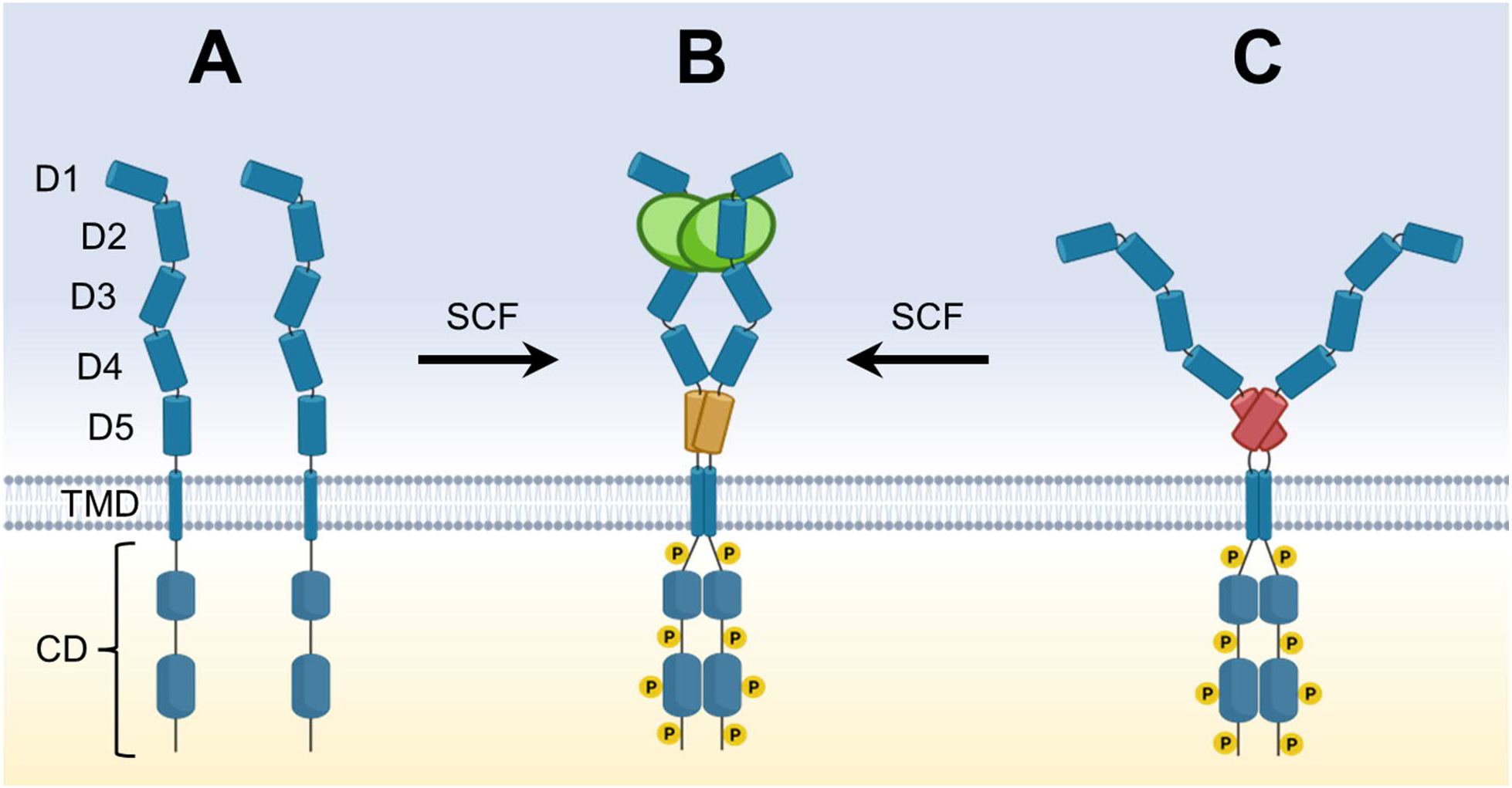
Structural plasticity of the ligand-independent constitutively active oncogenic KIT mutant. Binding of SCF to monomeric wild-type KIT (*A*) results in the formation of KIT:SCF dimers (*B*). In KIT:SCF dimers, the ECDs are held together by SCF binding to D1–D3, by D4:D4’ homotypic interactions via symmetric contacts, and by D5:D5’ interactions mediated by an asymmetric interface. The ECDs of ligand-independent, constitutively active KIT(T417I,Δ418-419) dimers adopt a V-shaped conformation solely held together by D5:D5’ contacts (*C*). The D5:D5’ complex of this constitutively active oncogenic KIT mutant adopts a strongly tilted, symmetric conformation with sufficient affinity to trigger KIT dimerization and activation in the absence of SCF binding. Binding of SCF to KIT(T417I,Δ418-419) dimers restores a wild-type-like conformation of the ECDs with an asymmetric D5:D5’ interface (*B*). CD, cytoplasmic domain; TMD, transmembrane domain.

We have previously explored and compared the response of six different KIT oncogenic mutants—including the two mutants described in this manuscript—to the FDA approved KIT tyrosine kinase inhibitors imatinib and sunitinib, to therapeutic antibodies targeting D4 of KIT, and to toxin-conjugated anti-KIT antibodies (22). We concluded that “each of the six major KIT oncogenic mutants exhibits distinct properties and responds differently to targeted therapies” that include single agents or combination of two treatments. The remarkable structural plasticity of the oncogenic KIT(T417I,Δ418-419) mutant described in the present manuscript may offer new therapeutic regimen for treatment of such tumors. It was previously reported that oncogenic KIT mutants that act inside the cell can be flushed out to the cell membrane by interfering with their tyrosine kinase activities using tyrosine kinase inhibitors such as imatinib and sunitinib (23, 24). We surmise that once relocated to the cell membrane (after imatinib or sunitinib treatment), an oncogenic mutant such as KIT(T417I,Δ418-419) that is treated with SCF could induce restoration of a wild-type-like KIT conformation to tame the harmful excessive activity of the oncogenic mutant. A combination therapy involving a tyrosine kinase inhibitor together with ligand treatment may take advantage of the structural plasticity of certain oncogenic mutations for therapeutic purposes.

## Materials and Methods

### Expression and purification of SCF

For cryo-EM and cell autophosphorylation studies, SCF was expressed in *E*.*coli* BL21-CodonPlus (DE3)-RIPL, refolded from inclusion bodies, and purified as described previously (25).

### Expression and purification of full-length KIT for cryo-EM studies

For structural studies, a kinase-inactive KIT (K623A) mutant was introduced into full-length wild-type and mutants of KIT to increase yield and homogeneity. The cDNA encoding KIT (Uniprot: P10721-2, residues 32– 976) with a N-terminal FLAG-tag and a C-terminal 6×His-tag preceded by a two-residues linker were subcloned into pFastBac 1 vector (Gibco). Sequence transposition into bacmid DNA using DH10Bac cells (Gibco) and analysis of the recombinant bacmid were performed following the manufacturer’s instructions (ExpiSf Expression System, Gibco). ExpiSf9 cells were cultured in ExpiSf9 CD medium to a density of 5×10^6^ cells/mL, and infected with P0 baculovirus stock. Infected cells were incubated for 48 h at 27 °C. Cells were harvested by centrifugation (4000×g, 10 min), resuspended in lysis buffer (10% glycerol, 20 mM HEPES pH 7.4, 200 NaCl, cOmplete protease inhibitor mixture (Roche), 1 mg/mL DNAse I (Roche)), and lysed by pressure homogenization using an Emulsiflex C3 cell disruptor (Avestin) at 5,000 psi. The lysate was centrifuged at 700×g for 30 min, and the supernatant was subjected to ultracentrifugation (Beckman MLA-80, 55,000 rpm, 45 min). The pellet was suspended in resuspension buffer (5% glycerol, 20 mM HEPES pH 7.4, 200 NaCl, cOmplete protease inhibitor mixture (Roche)) at a concentration of 4 mg/mL using a tissue grinder. For the formation of KIT:SCF complexes, SCF was added at a concentration of 2 μM. After incubation over night while rotating head-over-head at 4°C, 1% DDM (w/w) was added, followed by a further 1 h of incubation. The sample was ultracentrifuged (Beckman 45Ti, 40,000 rpm, 1 h), and the supernatant was applied to an anti-FLAG M2 affinity gel (Sigma). The resin was washed with resuspension buffer additionally containing 0.04% DDM, and the sample was eluted using 3×FLAG peptide (50 μM). Protein-containing fractions were combined, and amphipol A8-35 (Anatrace) was added at a ratio of 1:4 (w/w), and the sample was incubated for 30 min at 4°C using a rocking shaker. Subsequently, Biobeads SM-2 resin (Bio-Rad) was added at a ratio of 1:5 (w/v), and the sample was incubated for 2 h at 4°C using a rocking shaker. After removal of the Biobeads SM-2 resin, the sample was subjected to glycerol gradient ultracentrifugation in the presence of a chemical fixation reagent (5– 20% glycerol, 0–0.2% glutaraldehyde, 20 mM HEPES pH 7.4, 200 mM NaCl) (19). Gradients were prepared using a Gradient Master 108 (BioComp Instruments). Samples were ultracentrifuged using a Beckman SW41Ti rotor at 40,000 rpm for 18 h at 4°C. Samples were fractionated into 500 μL fractions from top to bottom, and glutaraldehyde was quenched by adding 100 mM Tris pH 7.4. Individual fractions were analyzed by SDS-PAGE using Pierce Silver Stain (Thermo Fisher Scientific). Fractions containing pure KIT or KIT:SCF dimers were gently concentrated using a 100 kDa MWCO concentrator (GE Healthcare) at 100×g to a concentration of 6.5 mg/mL, and flash-frozen in liquid nitrogen.

### Negative staining EM sample preparation

Prior to using a sample for cryo-EM, the quality of the sample was confirmed by negative staining EM. Protein samples (4 μL, 0.05 mg/mL) were applied on the carbon side of glow-discharged carbon-coated holey copper grids (EMS CF300-CU). After 1 min, excess protein solution was wicked away using filter paper. Particles were stained by floating the grid on a 50 μL drop of freshly prepared 1.5% (w/v) uranyl formate solution for 1 min, followed by wicking away excess liquid with filter paper. The procedure of staining and wicking away excess liquid was repeated once, followed by air-drying of the grid. Grids were screened on a FEI Tecnai T12 TEM operated at 120 kV, equipped with a Gatan UltraScan 4000 (4k×4k) CCD camera. 2D classification was performed using RELION 3.1 (26).

### Cryo-EM sample preparation

Samples were used at concentrations of 5.5–6.0 mg/mL in sample buffer, to which fluorinated FC-8 was added at a final concentration of 0.1% (v/v). All samples were prepared with glow-discharged holey carbon grids (C-flat 1.2/1.3, 300 mesh, gold support). Samples were vitrified using a Gatan CP3 (no wait time, room temperature, approximately 90% relative humidity, 3–4 s blotting time, blot force 0), and plunge-frozen in liquid ethane cooled by liquid nitrogen. Grids were screened on a Glacios Cryo-TEM (Thermo Fisher Scientific) operated at 200 keV, equipped with a K2 summit direct detection camera.

### Cryo-EM dataset collection

Datasets of wild-type KIT:SCF dimers, KIT(DupA502,Y503):SCF dimers, and KIT(T417I,Δ418-419) dimers were collected on a FEI Titan Krios G2 300 keV TEM, equipped with a Gatan Quantum LS imaging filter and a Gatan K3 direct electron detector, at the Yale Cryo-EM resources. SerialEM was used for automatic data acquisition (27). The dataset of KIT(T417I,Δ418-419):SCF dimers was collected on a FEI Titan Krios G3i 300 keV TEM, equipped with a BioQuantum energy filter and a Gatan K3 direct electron detector, at the LBMS located at the Brookhaven National Laboratory. EPU (Thermo Fisher Scientific) was used for automatic data acquisition. All datasets were collected in super-resolution counting mode, operated in correlated-double sampling (CDS) imaging mode. Data collection parameters are detailed in Table S1.

### Cryo-EM data processing

All datasets of wild-type and mutants of KIT in complex with SCF were processed according to the below described workflow. Individual data processing flow charts with further details are shown in Fig. S1B for wild-type KIT:SCF, Fig. S5B for KIT(DupA502,Y503):SCF, and Fig. S8C for KIT(T417I,Δ418-419):SCF. The dataset of wild-type KIT:SCF was processed using cryoSPARC 3.1.0, and datasets of KIT(DupA502,Y503):SCF as well as KIT(T417I,Δ418-419):SCF were processed using cryoSPARC 3.2.0. Raw movies were corrected for beam-induced motion using patch motion correction (2×binning). CTF parameters were estimated using patch CTF estimation (default settings). Micrographs of poor quality, e.g., with large ice contaminants or cracked vitreous ice, were manually removed. Initial particles were picked from a small fraction of micrographs using reference-free blob picking and subjected to 2D classification. Well-resolved 2D class averages showing different projections of KIT molecules were used as templates for template-based particle picking. Particle picks were extracted, and initially cleaned from incorrect picks (e.g., contaminants) by multiple rounds of 2D classification (default settings, 50–200 classes). Particles were further cleaned by performing two iterative rounds of a six-classes ab-initio reconstruction followed by heterogeneous refinement using the six ab-initio reconstructions as input models, followed by removal of ‘junk’ 3D classes. Subsequently, particles from clearly defined 3D classes were combined and used for the generation of a one-class ab-initio reconstruction, followed by re-centering of the particle by re-extraction using aligned shifts, CTF refinement, and non-uniform (NU) refinement (28). Generally, NU refinement of all selected particles together resulted in a map with better resolution and more pronounced features compared to 3D classification using cryoSPARC or RELION and subsequent refinement of the individual classes or combinations thereof. Following NU refinement, a soft mask covering the entire ECD was created, and used for non-uniform local refinement of the ECD. All refinements were performed in C1 symmetry to account for the asymmetric conformation of D5:D5’. Sharpened and unsharpened maps were obtained from cryoSPARC. In the case of wild-type KIT:SCF, the map was post-processed using deepEMhancer 0.13 using the default tightTarget model (29). The continuous flexibility of wild-type KIT:SCF (Fig. S1G) was analyzed using cryoSPARC’s 3D variability analysis (3DVA) algorithm (21).

The dataset of KIT(T417I,Δ418-419) dimers was processed using cryoSPARC 3.1.0 following a similar workflow as described above, and as detailed in Fig. S6C. Homogeneous refinement using cryoSPARC resulted in a map well-defined for the region of D5:D5’, but otherwise with incompletely defined ECDs. Thus, particles were exported to RELION 3.1, and subjected to 3D classification into six classes (Fig. S6D). The resulting class with the best-defined ECD was used for the creation of a loose soft mask covering the entire ECD, excluding the transmembrane and cytoplasmic regions. Using this mask, a masked 3D classification of the ECD into six classes was performed. Subsequently, particles of classes with the best resolved ECD were combined and subjected to masked 3D refinement of the ECD, generating a map where both ECDs are fully defined (Fig. 6D).

All map resolutions are reported for the gold-standard FSC threshold criterion of 0.143, calculated from both half maps using cryoSPARC or RELION. Local resolution maps were calculated using cryoSPARC.

### Model building and refinement

As starting model for the cryo-EM model of the ECD of wild-type KIT:SCF, the crystal structure of the truncated ECD of KIT:SCF (PDB ID 2E9W) was docked into the cryo-EM map using UCSF Chimera. The model was manually built and adjusted using Coot (version 0.96) (30) and ISOLDE (version 1.0b3) (31), using the unsharpened and the deepEMhancer post-processed maps. The deepEMhancer post-processed map appears discontinuous for C-terminal residues 507–509 of protomer A and 508–510 of protomer B. Therefore, these residues were modeled using the unsharpened map. Despite the lower resolution of D5:D5’ compared to the rest of the ECD (Fig. S1B), interface residues were readily identified and modeled due to the bulky side chains of the interface residues (Y418, Y503, F504, F506) resulting in characteristic and clear side chain features in the cryo-EM map (Fig. S1F). Loops of D5 and D5’ proximal to the membrane, and β-strands βD and βD’ (Fig. S2B) were omitted from the model due to insufficiently defined cryo-EM density preventing confident modelling. Side chains of residues with insufficiently defined cryo-EM density were removed from the model. Following model building, the model was refined using real-space refinement in Phenix (version 1.02.1-4487-000) (32). Model quality was validated using MolProbity (33). The models of KIT(DupA502,Y503):SCF and KIT(T417I,Δ418-419):SCF were built using the final cryo-EM model of wild-type KIT:SCF as starting model. Model building and refinement was performed similarly as described for wild-type KIT:SCF. Adjustments of the models were performed using the sharpened and unsharpened maps. Loops of D5 and D5’ proximal to the membrane, and β-strands βD and βD’ were omitted from both models. Due to the low resolution of D5:D5’ in KIT(T417I,Δ418-419):SCF (Fig. S8C), mostly only the main chains were modeled. Model refinement and validation statistics are listed in Table S1. Software was curated by SBGRID (34).

Structures were visualized and figures were prepared using UCSF Chimera (35) and ChimeraX (36). Buried surface areas were calculated using the PISA server from the European Bioinformatics Institute (37). The structure-based sequence alignments (Fig. 3A) were generated by superposing β-sheets βC-βF-βG of D5 domains. Due to a lack of high-resolution structural data for D5, AlphaFold (38) structure predictions were used for PDGFR*α* (UniProt P16234, AF-P16234-F1-model_v2), PDGFR*β* (UniProt P09619, AF-P09619-F1-model_v2), CSF1R (UniProt P07333, AF-P07333-F1-model_v2), and FLT3 (UniProt P36888, AF-P36888-F1-model_v2).

### Cloning of constructs for cell experiments

By using restriction sites EcoRI and ApaI, KIT DNA inserts containing the desired mutations were generated by PCR amplification using the Phusion High-Fidelity DNA Polymerase (New England Biolabs). The inserts were subcloned into either pFastBac 1 vector for structural studies or pBABE-puro vector for cell-based studies. A cDNA encoding for full-length human PDGFR*β* (NP_002600.1) was amplified by PCR and subcloned into lentiviral transfer plasmid pLenti CMV Hygro DEST. Using AgeI and MfeI restriction sites, PDGFR*β* DNA inserts containing the desired mutations were generated by PCR amplification using the Phusion High-Fidelity DNA Polymerase (New England Biolabs) and subcloned into pLenti CMV Hygro DEST for cellular studies. Recombinant plasmids were confirmed by restriction enzyme digestion and by DNA sequencing (Keck DNA Sequencing Facility at Yale). Primers used for the generation of the inserts are listed in Table S2.

### Cell culture, immunoprecipitation, and immunoblotting experiments

As previously described (17), the retroviral pBABE-puro vector was used to generate NIH 3T3 cells stably expressing wild-type and mutants of KIT (1–972). MEFs deficient in endogenous PDGFR*α* and PDGFR*β* (16) were used to stably express wild-type and mutants of PDGFR*β* (1–1106) with a C-terminal HA-tag. The generation of lentivirus for expressing the various PDGFR*β* constructs was generated as previously described (39).

NIH 3T3 cells stably expressing wild-type and KIT mutants were culture at 37°C, 5% CO_2_ in DMEM (Gibco) supplemented with 5% FBS (Gibco), 5% BS (Gibco), 1% penicillin-streptomycin (Gibco), and 0.0001% puromycin (Gibco). Cells that reached 90% confluency were stimulated with SCF at increasing concentrations for 10 min at 37°C (Fig. 3B). MEFs stably expressing wild-type and PDGFR*β* mutants were culture at 37°C, 5% CO_2_ in DMEM (Gibco) supplemented with 10% FBS (Gibco), 1% penicillin-streptomycin (Gibco), and 0.015% hygromycin B (Gibco). Cells that reached 90% confluency were stimulated with PDGFR-BB (Sigma) at increasing concentrations for 10 min at 37°C (Fig. 3C). Following stimulation, NIH 3T3 cells and MEFs were washed twice with ice-cold PBS and lysed with 600 μL of lysis buffer (50 mM HEPES pH 7.5, 150 mM NaCl, 1.5 mM MgCl_2_, 1 mM EDTA, 1 mM EGTA, 10% glycerol, 25 mM NaF, 1 mM Na_3_VO_4_, 1% Triton-X 100, cOmplet protease inhibitor mixture (Roche)). Cell lysates were then clarified by centrifugation (16,000×g, 20 min, 4°C). KIT was immunoprecipitated from the supernatant with 6 μg monoclonal anti-KIT antibody (Santa Cruz Biotechnology, no. sc-13508) together with 30 μL protein G PLUS-agarose beads (Santa Cruz Biotechnology, no. sc-2002). The lysates were incubated overnight at 4°C using a rocking shaker. PDGFR*β* was immunoprecipitated from the supernatant with 25 μL of monoclonal anti-HA antibody conjugated to Sepharose beads (Cell Signaling, no. C29F4). KIT and PDGFR*β* immunocomplexes were washed three times with washing buffer (50 mM HEPES pH 7.5, 150 mM NaCl, 1.5 mM MgCl_2_, 1 mM EDTA, 1 mM EGTA, 10% glycerol, 25 mM NaF, 1 mM Na_3_VO_4_, 0.1% Triton-X 100, cOmplete protease inhibitor mixture (Roche)), and resuspended in 80 μL of reducing Laemmli buffer. Samples were separated on a 4–15% gradient SDS-PAGE gel, transferred to nitrocellulose membrane (Thermo Fisher Scientific), and immunoblotted with either anti-KIT antibody (Cell Signaling Technology, no. D3W6Y), anti-phospho KIT (Y703) antibody (Cell Signaling Technology, no. D12E12), anti-PDGFR*β* antibody (Cell Signaling Technology, no.3169S), or anti-phosphotyrosine (pTyr) antibody (Upstate Biotechnology, no. 4G10). All primary antibodies were used at a dilution of 1:1000. HRP-linked anti-rabbit IgG (Cell Signaling Technology, no. 7074S) or HRP-linked anti-mouse IgG (Cell Signaling Technology, no. 7076S) were used at a concentration of 1:2500. All blots were developed with enhanced chemiluminescent substrate (Bio-Rad Laboratories) and imaged using an iBright FL1000 device (Invitrogen). Blots were densitometrically quantified using the iBright Analysis Software.

## Data Availability

Cryo-EM maps and atomic model coordinates have been deposited in the EMDB and RCSB, respectively, under accession codes EMD-27408/8DFM, EMD-27410/8DFP, EMD-27411/8DFQ, EMD-27495, EMD-27496.

## Supplemental Information

Supplemental Fig. S1–S8, and Tables S1 and S2.

## Author contributions

S.G.K., N.B., I.L., and J.S. conceived project; N.B. and S.G.K. expressed and purified protein samples; S.G.K. and N.B. prepared cryo-EM samples with assistance from S.L. and W.M; N.B., S.G.K, and S.S. collected cryo-EM data under guidance of W.M.; S.G.K. and N.B. performed cryo-EM data processing and interpretation with assistance from S.L. and W.M.; N.B., S.G.K, and Y.S. designed mutagenesis studies and generated expression constructs; N.B., S.G.K., J.M., L.K., and I.L. performed cell signaling studies; S.G.K., N.B., and J.S. wrote the manuscript with editing from all authors; I.L. and J.S. supervised the project.

## Acknowledgements

We thank the members of the Schlessinger laboratory for valuable discussions, and Seong An and Nathan Kucera for proof-reading of the manuscript. We thank the Center for Cellular and Molecular Imaging, Electron Microscopy Facility at Yale Medical School, Science Hill, and West campus for assistance with the work presented here. In particular, we thank Marc C. Llaguno, Shenping Wu, and Jianfeng Lin for assistance during data collection. The Laboratory for BioMolecular Structure (LBMS) is supported by the DOE Office of Biological and Environmental Research (KP160711).

## References

1. M. A. Lemmon, J. Schlessinger, Cell signaling by receptor tyrosine kinases. Cell 141, 1117–34 (2010).

2. L. Z. Mi, et al., Simultaneous visualization of the extracellular and cytoplasmic domains of the epidermal growth factor receptor. Nat. Struct. Mol. Biol. 18, 984–989 (2011).

3. Y. Opatowsky, et al., Structure, domain organization, and different conformational states of stem cell factor-induced intact KIT dimers. Proc. Natl. Acad. Sci. U. S. A. 111, 1772–1777 (2014).

4. P. H. Chen, V. Unger, X. He, Structure of Full-Length Human PDGFRβ Bound to Its Activating Ligand PDGF-B as Determined by Negative-Stain Electron Microscopy. J. Mol. Biol. 427, 3921–3934 (2015).

5. E. Uchikawa, E. Choi, G. Shang, H. Yu, B. Xiao-Chen, Activation mechanism of the insulin receptor revealed by cryo-EM structure of the fully liganded receptor-ligand complex. Elife 8, 1–23 (2019).

6. Y. Huang, et al., A molecular mechanism for the generation of ligand-dependent differential outputs by the epidermal growth factor receptor. Elife 10, 1–32 (2021).

7. D. Diwanji, et al., Structures of the HER2–HER3–NRG1β complex reveal a dynamic dimer interface. Nature 600, 339–343 (2021).

8. L. K. Ashman, R. Griffith, Therapeutic targeting of c-KIT in cancer. Expert Opin. Investig. Drugs 22, 103–115 (2013).

9. R. Roskoski, The role of small molecule Kit protein-tyrosine kinase inhibitors in the treatment of neoplastic disorders. Pharmacol. Res. 133, 35–52 (2018).

10. C. L. Corless, C. M. Barnett, M. C. Heinrich, Gastrointestinal stromal tumours: Origin and molecular oncology. Nat. Rev. Cancer 11, 865–878 (2011).

11. J. Lennartsson, L. Rönnstrand, Stem cell factor receptor/c-Kit: From basic Science to clinical implications. Physiol. Rev. 92, 1619–1649 (2012).

12. A. Ullrich, J. Schlessinger, Signal Transduction by Receptors with Tyrosine Kinase Activity. Cell 61, 203–212 (1990).

13. M. A. Lemmon, D. Pinchasi, M. Zhou, I. Lax, J. Schlessinger, Kit receptor dimerization is driven by bivalent binding of stem cell factor. J. Biol. Chem. 272, 6311–6317 (1997).

14. H. Liu, X. Chen, P. J. Focia, X. He, Structural basis for stem cell factor-KIT signaling and activation of class III receptor tyrosine kinases. EMBO J. 26, 891–901 (2007).

15. A. V. Reshetnyak, et al., The strength and cooperativity of KIT ectodomain contacts determine normal ligand-dependent stimulation or oncogenic activation in cancer. Mol. Cell 57, 191–201 (2015).

16. Y. Yang, S. Yuzawa, J. Schlessinger, Contacts between membrane proximal regions of the PDGF receptor ectodomain are required for receptor activation but not for receptor dimerization. Proc. Natl. Acad. Sci. U. S. A. 105, 7681–7686 (2008).

17. S. Yuzawa, et al., Structural Basis for Activation of the Receptor Tyrosine Kinase KIT by Stem Cell Factor. Cell 130, 323–334 (2007).

18. K. Verstraete, S. N. Savvides, Extracellular assembly and activation principles of oncogenic class III receptor tyrosine kinases. Nat. Rev. Cancer 12, 753–766 (2012).

19. B. Kastner, et al., GraFix: Sample preparation for single-particle electron cryomicroscopy. Nat. Methods 5, 53–55 (2008).

20. J. Li, E. Choi, H. Yu, X. chen Bai, Structural basis of the activation of type 1 insulin-like growth factor receptor. Nat. Commun. 10, 1–11 (2019).

21. A. Punjani, D. J. Fleet, 3D variability analysis: Resolving continuous flexibility and discrete heterogeneity from single particle cryo-EM. J. Struct. Biol. 213, 107702 (2021).

22. X. Shi, et al., Distinct cellular properties of oncogenic KIT receptor tyrosine kinase mutants enable alternative courses of cancer cell inhibition. Proc. Natl. Acad. Sci. U. S. A. 113, E4784–E4793 (2016).

23. H. Bougherara, et al., The aberrant localization of oncogenic kit tyrosine kinase receptor mutants is reversed on specific inhibitory treatment. Mol. Cancer Res. 7, 1525–1533 (2009).

24. S. Tabone-Eglinger, et al., KIT mutations induce intracellular retention and activation of an immature form of the KIT protein in gastrointestinal stromal tumors. Clin. Cancer Res. 14, 2285–2294 (2008).

25. Z. Zhang, R. Zhang, A. Joachimiak, J. Schlessinger, X. P. Kong, Crystal structure of human stem cell factor: Implication for stem cell factor receptor dimerization and activation. Proc. Natl. Acad. Sci. U. S. A. 97, 7732–7737 (2000).

26. J. Zivanov, et al., New tools for automated high-resolution cryo-EM structure determination in RELION-3. Elife 7, 1–22 (2018).

27. D. N. Mastronarde, Automated electron microscope tomography using robust prediction of specimen movements. J. Struct. Biol. 152, 36–51 (2005).

28. A. Punjani, H. Zhang, D. J. Fleet, Non-uniform refinement: adaptive regularization improves single-particle cryo-EM reconstruction. Nat. Methods 17, 1214–1221 (2020).

29. R. Sanchez-Garcia, et al., DeepEMhancer: a deep learning solution for cryo-EM volume post-processing. Commun. Biol. 4, 1–8 (2021).

30. P. Emsley, K. Cowtan, Coot: model-building tools for molecular graphics. Acta Crystallogr., Sect. D Biol. Crystallogr. 60, 2126–2132 (2004).

31. T. I. Croll, ISOLDE: A physically realistic environment for model building into low-resolution electron-density maps. Acta Crystallogr. Sect. D Struct. Biol. 74, 519–530 (2018).

32. P. D. Adams, et al., PHENIX: A comprehensive Python-based system for macromolecular structure solution. Acta Crystallogr. Sect. D Biol. Crystallogr. 66, 213–221 (2010).

33. C. J. Williams, et al., MolProbity: More and better reference data for improved all-atom structure validation. Protein Sci. 27, 293–315 (2018).

34. A. Morin, et al., Collaboration gets the most out of software. Elife 2013, 1–6 (2013).

35. E. F. Pettersen, et al., UCSF Chimera - A visualization system for exploratory research and analysis. J. Comput. Chem. 25, 1605–1612 (2004).

36. E. F. Pettersen, et al., UCSF ChimeraX: Structure visualization for researchers, educators, and developers. Protein Sci. 30, 70–82 (2021).

37. E. Krissinel, K. Henrick, Inference of Macromolecular Assemblies from Crystalline State. J. Mol. Biol. 372, 774–797 (2007).

38. J. Jumper, et al., Highly accurate protein structure prediction with AlphaFold. Nature 596, 583–589 (2021).

39. E. Campeau, et al., A versatile viral system for expression and depletion of proteins in mammalian cells. PLoS One 4, e6529 (2009).

