## Supplemental Information for "Cryo-EM analyses of wild-type and oncogenic KIT mutants reveal structural oncogenic plasticity and a novel “Achilles heel” for therapeutic intervention"

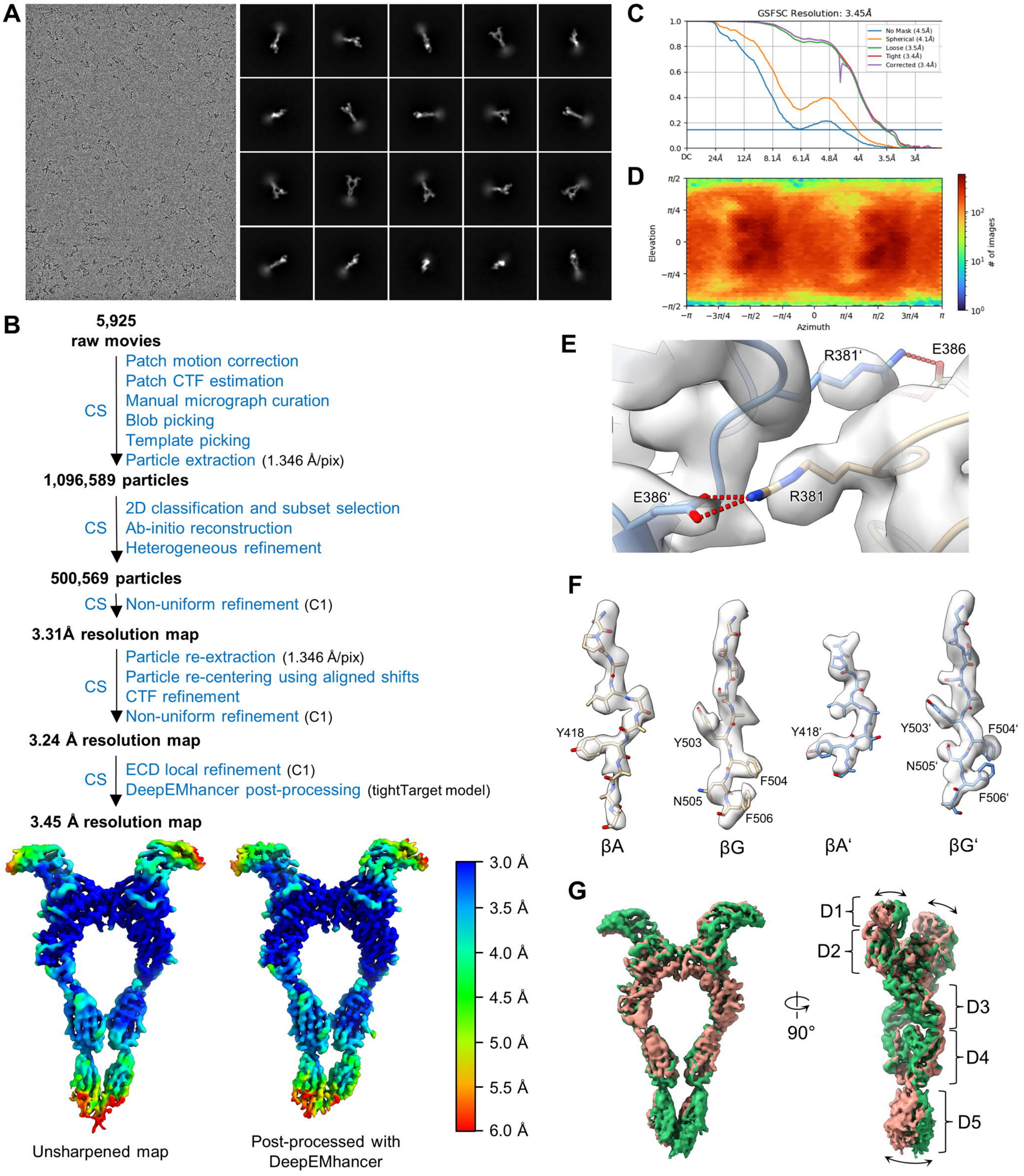


**Fig. S1.** Cryo-EM data processing and structural analysis of wild-type KIT:SCF dimers. (*A*) Representative cryo-EM micrograph and 2D class averages. (*B*) Cryo-EM data processing flow-chart. The local resolution map of the ECD local refinement (overall resolution: 3.45 Å) is displayed unsharpened (left) and post-processed using deepEMhancer (right). CS, cryoSPARC. (*C*) FSC curves of the ECD local refinement. (*D*) Angular distribution plot of particles used for ECD local refinement. (*E*) Cryo-EM density of the homotypic D4:D4’ salt bridge. (*F*) Cryo-EM density of residues of β-strands βA, βG, βA’, and βG’ forming the D5:D5’ interface. (*G*) Conformational flexibility analysis of the ECD using 3DVA of cryoSPARC. The two maps in red and green show the two most distant conformations of the motion solved for the first eigenvector. Significant motion (indicated by black arrows) is observed for domains D1, D1’, and for D5:D5’.

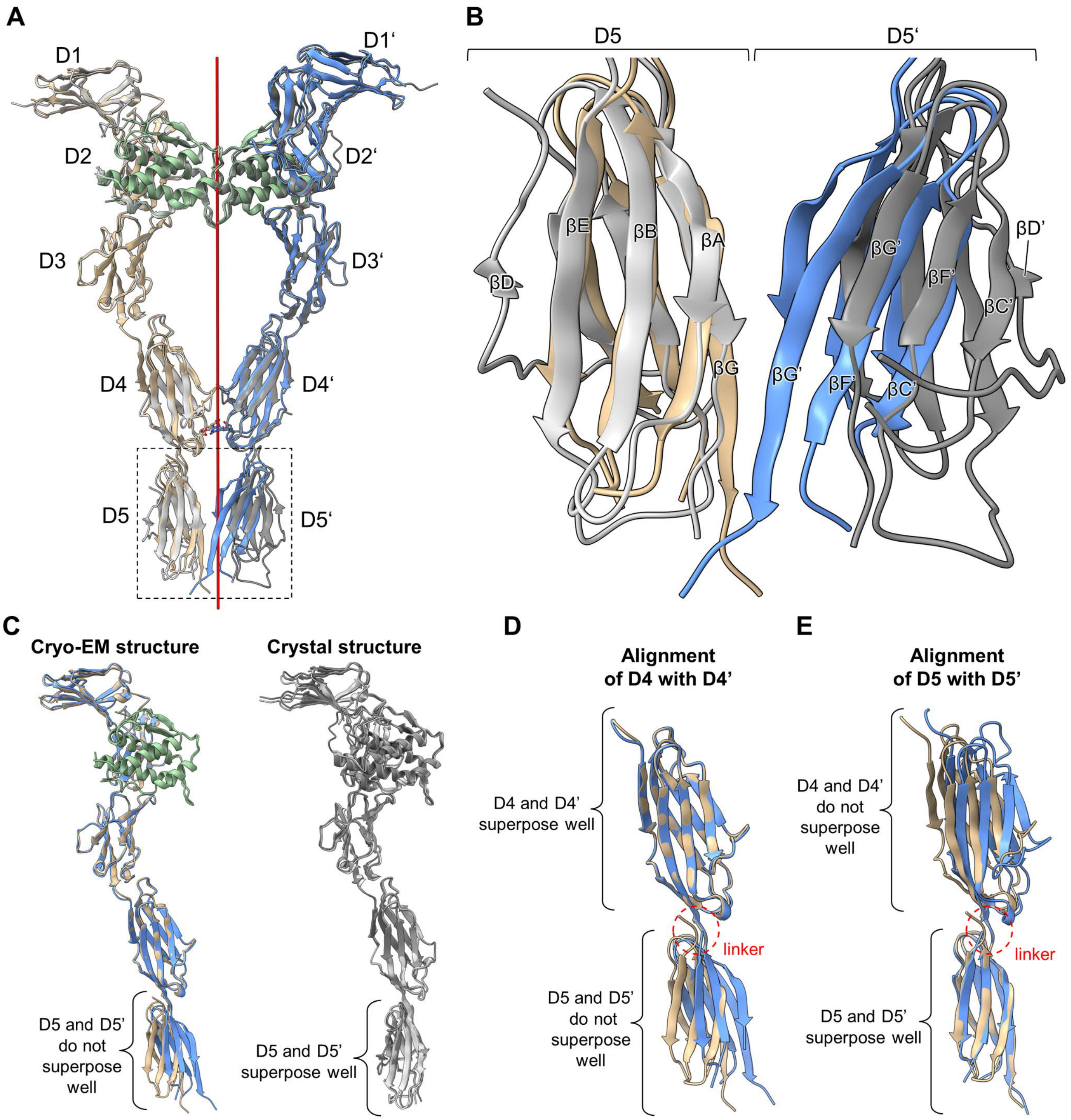


**Fig. S2.** Structural analysis of the asymmetric D5:D5' conformation in the cryo-EM structure of wild-type KIT:SCF dimers. (*A*) Superposition of the ECDs of the full-length cryo-EM structure (protomer A: beige; protomer B: blue; SCF homodimer: green) and the truncated crystal structure (PDB ID 2E9W; protomers A, B, and SCF in gray) of wild-type KIT:SCF dimers. The same color-code is used in *A*–*E*. The red line indicates the C2 symmetry axis. (*B*) Close-up view of the boxed D5:D5' superposition from *A*. β-strands βD and βD' were not included in the cryo-EM model due to poor local density in the cryo-EM map. (*C*) Superposition of KIT protomers A and B. As a result of the asymmetric conformation of D5:D5' in the cryo-EM structure, domains D5 and D5' do not superpose well (left). In contrast, the conformations of D5 and D5' in the crystal structure are symmetric, and thus superpose well (right). (*D*–*E*) Superpositions of regions D4D5 and D4'D5' of the cryo-EM structure. The tertiary conformations of D4, D4', D5, and D5' are largely conserved, and the asymmetric conformation of D5:D5' is enabled by the flexible linkers connecting D4 to D5 and D4' to D5'.

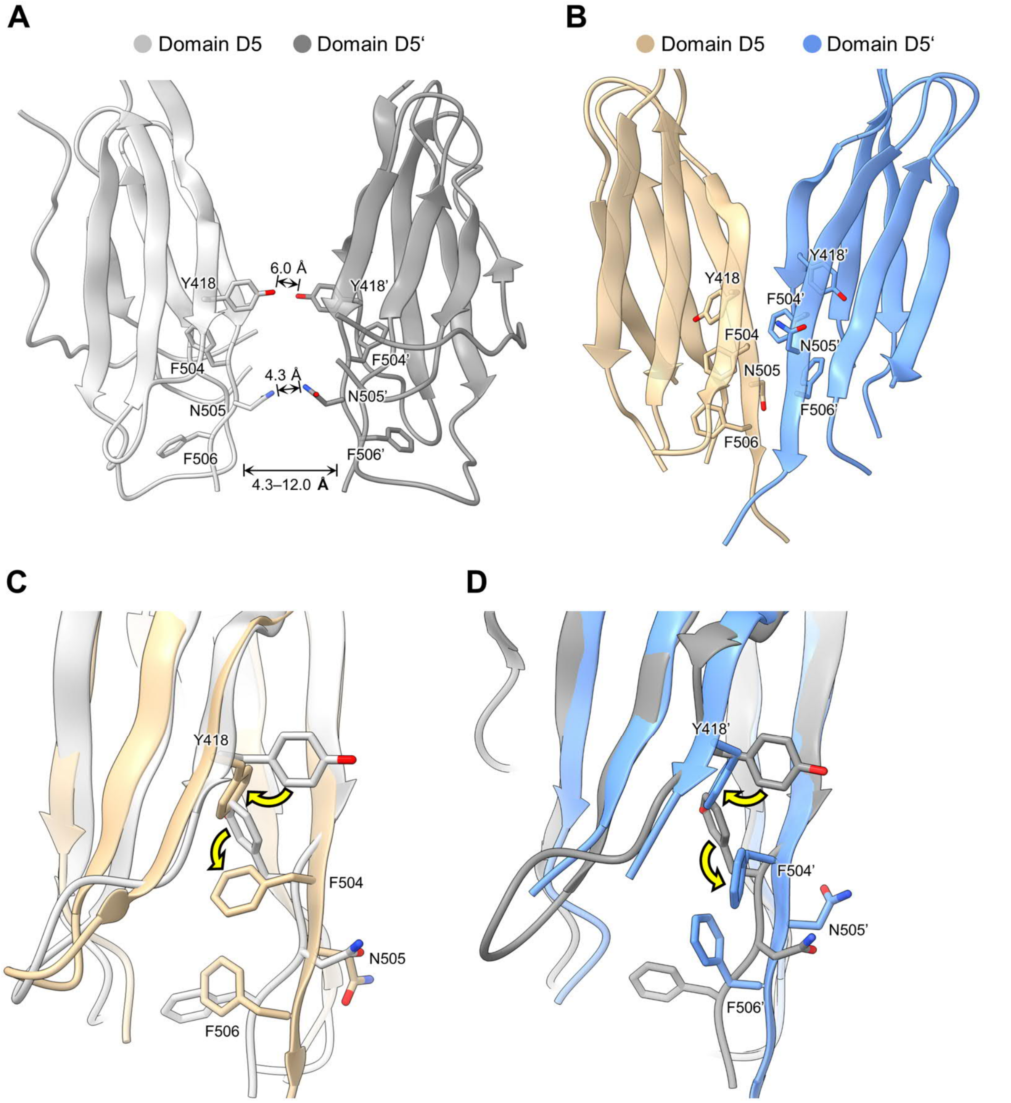


**Fig. S3.** D5:D5' contact formation requires conformational changes of interface residues. (*A*) Close-up view of domains D5 (light gray) and D5' (dark gray) of the crystal structure of the truncated ECD of wild-type KIT:SCF (PDB ID 2E9W). There is no direct contact between D5 and D5' (shortest distance: 4.3 Å, between N505 and N505'). (*B*) Close-up view of the D5:D5' complex of the cryo-EM structure of full-length wild-type KIT:SCF dimers. D5:D5' has an asymmetric quaternary conformation. (*C*) Superposition of D5 of the crystal structure (light gray) from panel *A* and D5 of the cryo-EM structure (beige) from panel *B*. The significant conformational changes of Y418 and F504 upon D5:D5' complexation are indicated by arrows. (*D*) Superposition of D5' of the crystal structure (dark gray) from panel *A* and D5' of the cryo-EM structure (blue) from panel *B*. The significant conformational changes of Y418' and F504' upon D5:D5' complexation are indicated by arrows.

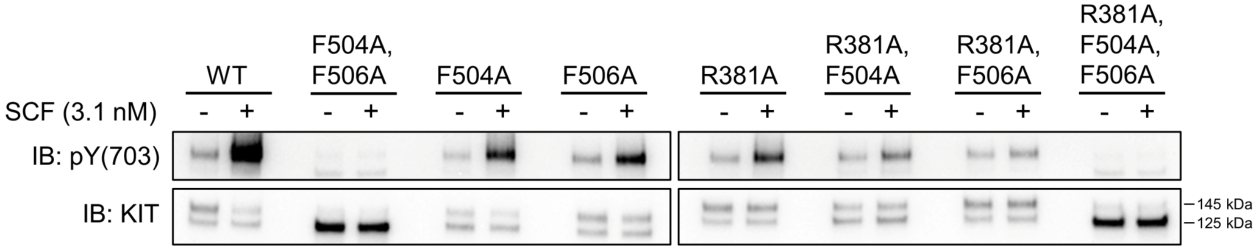


**Fig. S4.** Tyrosine autophosphorylation of wild-type and mutants of KIT after stimulation with SCF. SCF stimulated or unstimulated NIH 3T3 cells stably expressing WT KIT or KIT mutants were lysed and subjected to immunoprecipitation with anti-KIT antibody followed by SDS-PAGE and immunoblotting with anti-KIT or anti-phospho-KIT (Y703) antibodies. Full-length KIT migrates as two bands with an apparent MW of 145 kDa and 125 kDa. The 145 kDa band corresponds to mature and fully glycosylated KIT, whereas the 125 kDa band corresponds to immature and only partially glycosylated KIT. Only mature and fully glycosylated KIT is expressed on the cell surface, and subject to activation by SCF (1).

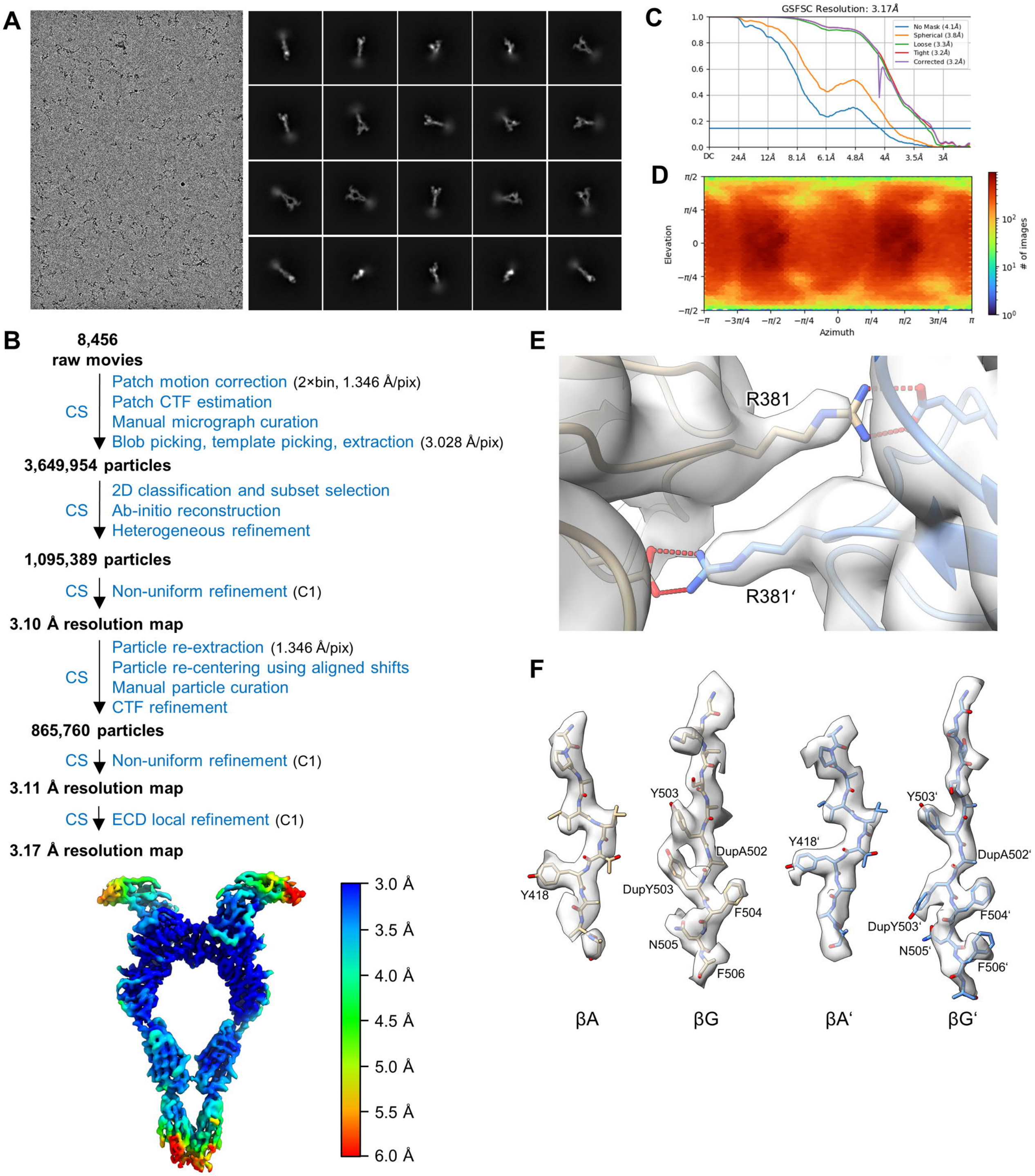


**Fig. S5.** Cryo-EM data processing and structural analysis of KIT(DupA502,Y503):SCF dimers. (*A*) Representative cryo-EM micrograph and 2D class averages. (*B*) Cryo-EM data processing flow-chart. The unsharpened local resolution map of the ECD local refinement (overall resolution: 3.17 Å) is displayed. CS, cryoSPARC. (*C*) FSC curves of the ECD local refinement. (*D*) Angular distribution plot of the particles used for the ECD local refinement. (*E*) Cryo-EM density of the homotypic D4:D4' salt bridge. (*F*) Cryo-EM density of residues of β-strands βA, βG, βA', and βG' forming the D5:D5' interface.

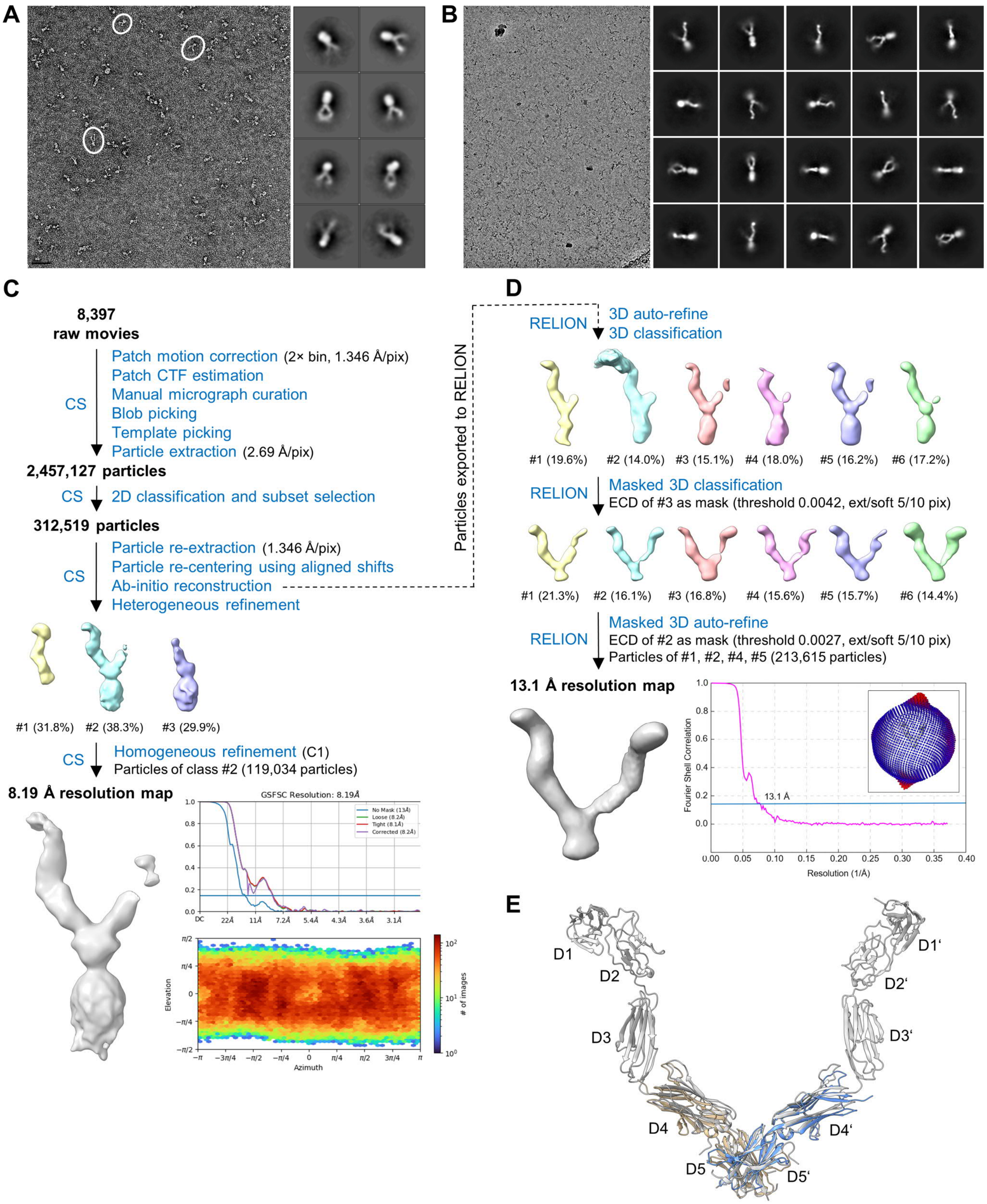


**Fig. S6.** Cryo-EM data processing and structural analysis of KIT(T417I,Δ418-419) dimers. (*A*) Representative negative staining EM micrograph and 2D class averages. (*B*) Representative cryo-EM micrograph and 2D class averages. (*C*) Cryo-EM data processing flow-chart of the homogeneous refinement of the entire complex (8.19 Å). CS, cryoSPARC. (*D*) Cryo-EM data processing flow-chart of the ECD masked refinement (13.1 Å, mask-uncorrected FSC). (*E*) Superposition of the crystal structure of D4D5 fragment dimers of KIT(T417I,Δ418-419) (PDB ID 4PGZ; protomer A in beige, protomer B in blue) and the cryo-EM structure of the ECD of full-length KIT(T417I,Δ418-419) dimers (in gray).

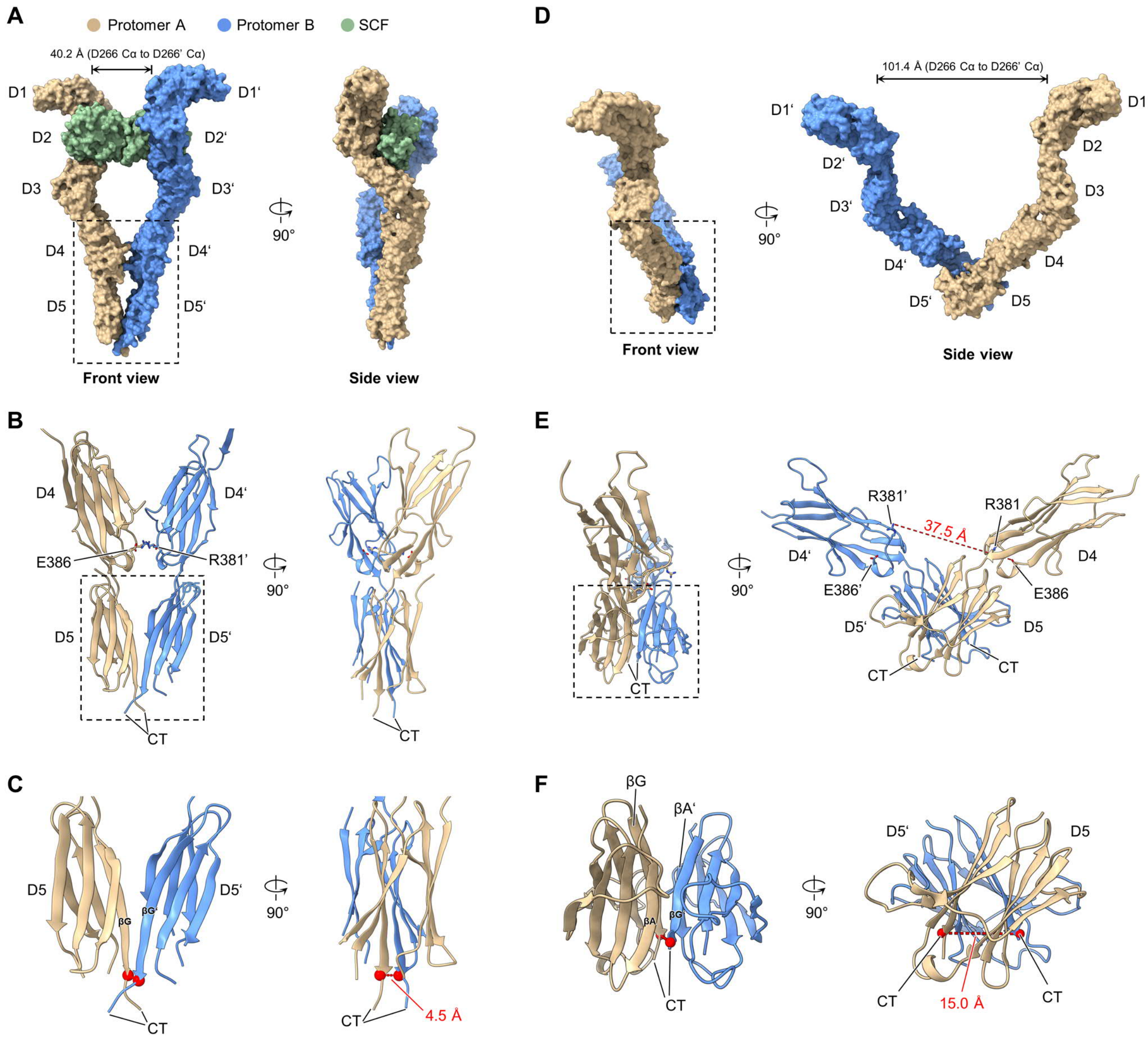


**Fig. S7.** Comparison of the ECD structures of full-length wild-type KIT:SCF dimers and full-length KIT(T417I,Δ418-419) dimers. (*A*) Space-filling model of the cryo-EM structure of wild-type KIT:SCF dimers from Fig. 2A. Protomer A in beige, protomer B in blue, and SCF in green. The same color-code is used in *A*–*F*. (*B*) Close-up view of the boxed D4D5:D4'D5' complex from *A*. CT, C-terminus. (*C*) Close-up view of the boxed D5:D5' complex from *B*. Cα atoms of C-terminal residues A507 and F508' are shown as red spheres. (*D*) Space-filling model of the cryo-EM structure of KIT(T417I,Δ418-419) dimers from Fig. 5C. (*E*) Crystal structure of D4D5 fragment dimers of KIT(T417I,Δ418-419) (PDB ID 4PGZ) corresponding to the boxed region from *D*. (*F*) Close-up view of the boxed D5:D5' complex from *E*. Cα atoms of C-terminal residues N505 and N505' are shown as red spheres.

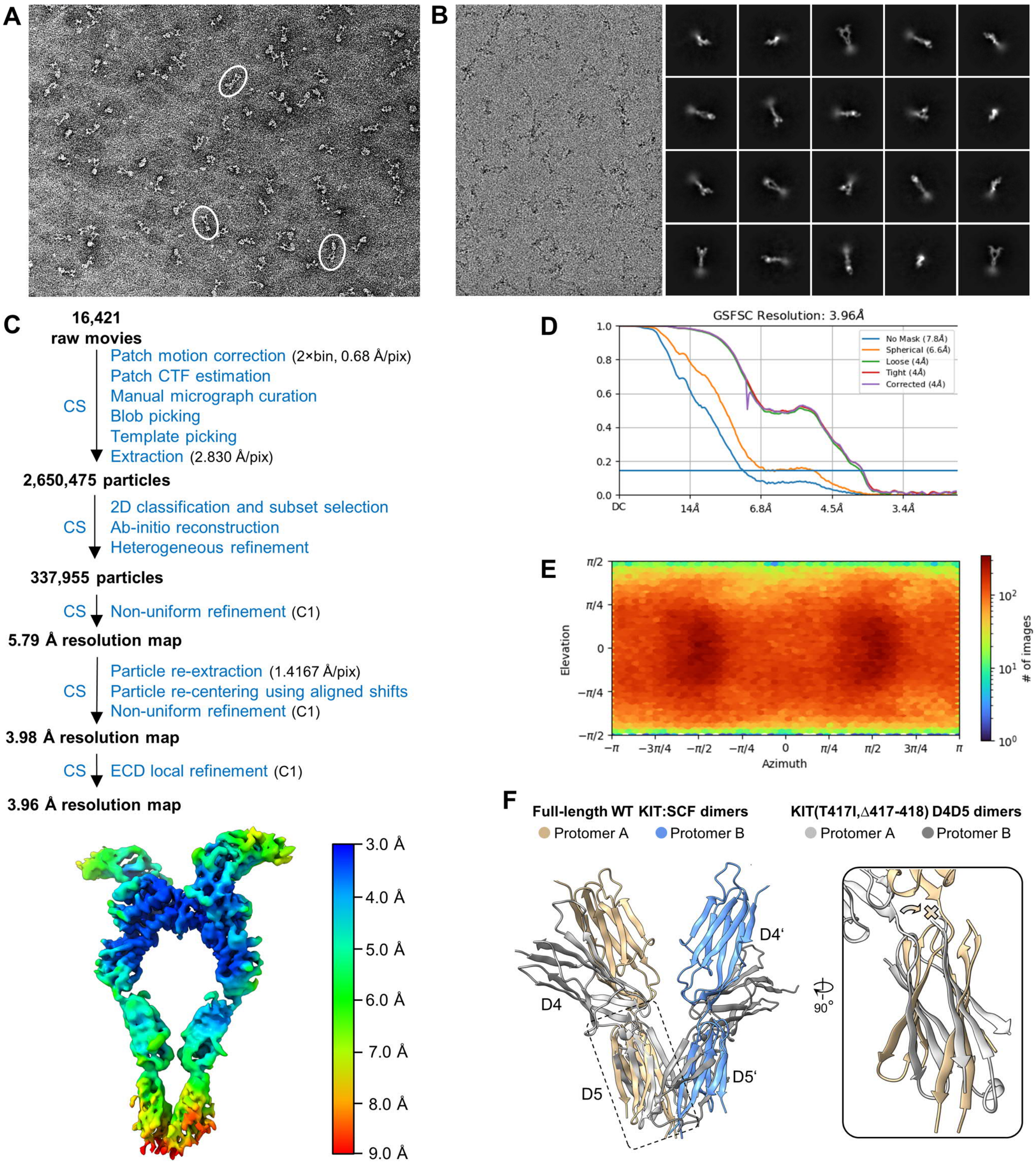


**Fig. S8.** Cryo-EM data processing and structural analysis of KIT(T417I,Δ418-419):SCF dimers. (*A*) Representative negative staining EM micrograph. Examples of single particles of KIT(T417I,Δ418-419):SCF dimers showing the characteristic shape of the ligand-bound dimeric receptor are encircled in white. (*B*) Representative cryo-EM micrograph and 2D class averages. (*C*) Cryo-EM data processing flow-chart. The unsharpened local resolution map of the ECD local refinement (overall resolution: 3.96 Å) is displayed. CS, cryoSPARC. (*D*) FSC curves of the ECD local refinement. (*E*) Angular distribution plot of the particles used for the ECD local refinement. (*F*) Superposition of the cryo-EM structure of full-length KIT(T417I,Δ418-419):SCF dimers and the crystal structure of D4D5 fragment dimers of KIT(T417I,Δ418-419). Due to steric pressure, binding of SCF to KIT(T417I,Δ418-419) dimers converts the tilted conformation of D5:D5' to the wild-type-like conformation of D5:D5' characterized by parallel-oriented domains.

**Table S1. Cryo-EM data collection, model refinement, and validation statistics**

| | **KIT:SCF dimers**<br>EMD-27408<br>PDB 8DFM | **KIT(DupA502, Y503):SCF dimers**<br>EMD-27410<br>PDB 8DFP | **KIT(T417I, Δ418-19) dimers**<br>EMD-27495 | **KIT(T417I, Δ418-19) dimers**<br>EMD-27496 | **KIT(T417I, Δ418-419):SCF dimers**<br>EMD-27411<br>PDB 8DFQ |
|---|---|---|---|---|---|
| **Data collection** | | | | | |
| Microscope | Titan Krios G2 | Titan Krios G2 | Titan Krios G2 | Titan Krios G2 | Titan Krios G3i |
| Detector | Gatan K3 with EF | Gatan K3 with EF | Gatan K3 with EF | Gatan K3 with EF | Gatan K3 with EF |
| Voltage (keV) | 300 | 300 | 300 | 300 | 300 |
| Magnification | ×64,000 | ×64,000 | ×64,000 | ×64,000 | ×130,000 |
| Pixel size (Å) | 1.346 | 1.346 | 1.346 | 1.346 | 0.680 |
| Exposure rate ($e^-$/pixel/s) | 10.02 | 10.5 | 10.537 | 10.537 | 15.0 |
| No. of frames per movie | 74 | 74 | 74 | 74 | 49 |
| Exposure time (s) | 8.9 | 11.134 | 11.134 | 11.134 | 1.52 |
| Total exposure ($e^-/Å^2$) | 49.23 | 64.52 | 64.76 | 64.76 | 49.31 |
| Defocus range (μm) | –1.0 to –2.5 | –1.2 to –2.2 | –1.3 to –2.3 | –1.3 to –2.3 | –0.5 to –1.5 |
| **Data processing** | | | | | |
| Number of movie stacks | 5,925 | 8,456 | 8,397 | 8,397 | 16,421 |
| Extraction box size (pixels) | 450 | 450 | 400 | 400 | 1,000 |
| Initial particle projections (no.) | 1,096,589 | 3,649,954 | 2,457,127 | 2,457,127 | 2,650,475 |
| Final particle projections (no.) | 500,569 | 865,760 | 119,034 | 213,615 | 337,955 |
| Map resolution (Å) | 3.45 | 3.17 | 8.19 | 13.1 | 3.96 |
| Symmetry imposed | C1 | C1 | C1 | C1 | C1 |
| FSC threshold | 0.143 | 0.143 | 0.143 | 0.143 | 0.143 |
| **Structure refinement** | | | | | |
| Initial model used | 2E9W | 8DFM | | | 8DFM |
| Model composition | | | | | |
| Nonhydrogen atoms | 8,400 | 8,421 | | | 7,719 |
| Protein residues | 1,126 | 1,118 | | | 1,114 |
| Ligand | 10 | 10 | | | 7 |
| B factors ($Å^2$) | | | | | |
| Protein | 117.0 | 119.4 | | | 145.3 |
| Ligand | 140.0 | 136.2 | | | 143.4 |
| R.m.s. deviations | | | | | |
| Bond lengths (Å) | 0.005 | 0.004 | | | 0.004 |
| Bond angles (°) | 1.1 | 1.1 | | | 1.1 |
| Validation | | | | | |
| MolProbity score | 0.86 | 0.82 | | | 0.78 |
| Clashscore | 1.31 | 1.11 | | | 0.91 |
| Poor rotamers (%) | 0 | 0 | | | 0.15 |
| Ramachandran plot: | | | | | |
| Favored (%) | 98.01 | 98.45 | | | 98.26 |
| Allowed (%) | 1.99 | 1.55 | | | 1.74 |
| Disallowed (%) | 0 | 0 | | | 0 |

**Table S2. Primers used in the study**

| Mutation | Direction | Nucleotide sequence of primer |
| --- | --- | --- |
| KIT R381A | Forward | 5’-GAACTTCATCTAACGGCATTAAAAGGCACCGAAGGAG-3’ |
| KIT R381A | Reverse | 5’-CTCCTTCGGTGCCTTTTAATGCCGTTAGATGAAGTTC-3’ |
| KIT F504A | Forward | 5’-CAAGACTTCTGCCTATGCTAACTTTGCATTTAAAGAGC-3’ |
| KIT F504A | Reverse | 5’-GCTCTTTAAATGCAAAGTTAGCATAGGCAGAAGTCTTG-3’ |
| KIT F506A | Forward | 5’-GACTTCTGCCTATTTTAACGCTGCATTTAAAGAGCAAATC-3’ |
| KIT F506A | Reverse | 5’-GATTTGCTCTTTAAATGCACGGTTAAAATAGGCAGAAG-3’ |
| KIT F504A,F506A | Forward | 5’-CTTCTGCCTATGCTAACGCTGCATTTAAAGAGCAAATC-3’ |
| KIT F504A,F506A | Reverse | 5’-GATTTGCTCTTTAAATGCACGGTTACGATAGGCAGAAG-3’ |
| KIT T417I,Δ418-419 | Forward | 5’-CAAAACCAGAAATCCTGATCAGGCTCGTGAATGGC-3’ |
| KIT T417I,Δ418-419 | Reverse | 5’-GCCATTCACGAGCCTGATCAGGATTTCTGGTTTTG-3’ |
| KIT DupA502,Y503 | Forward | 5’-GTGGGCAAGACTTCTGCCTATGCCTATTTTAACTTTGCATTTAAAG-3’ |
| KIT DupA502,Y503 | Reverse | 5’-CTTTAAATGCAAAGTTAAAATAGGCATAGGCAGAAGTCTTGCCCAC-3’ |
| PDGFR*β* V521A | Forward | 5’-CAGGACACGCAGGAGGCCATCGTGGTGCCACAC-3’ |
| PDGFR*β* V521A | Reverse | 5’-GTGTGGCACCACGATGGCCTCCTGCGTGTCCTG-3’ |
| PDGFR*β* V523A | Forward | 5’-CACGCAGGAGGTCATCGCGGTGCCACACTCCTTG-3’ |
| PDGFR*β* V523A | Reverse | 5’-CAAGGAGTGTGGCACCGCGATGACCTCCTGCGTG-3’ |
| PDGFR*β* R385A | Forward | 5’-GAGCTGACACTGGTTGCCGTGAAGGTGGCAGAG-3’ |
| PDGFR*β* R385A | Reverse | 5’-CTCTGCCACCTTCACGGCAACCAGTGTCAGCTC-3’ |

### Supplemental References


1. X. Shi, *et al.*, Distinct cellular properties of oncogenic KIT receptor tyrosine kinase mutants enable alternative courses of cancer cell inhibition. *Proc. Natl. Acad. Sci. U. S. A.* **113**, E4784–E4793 (2016).
